# Lineage-informed factor analysis reveals heritable programs of single-cell gene expression

**DOI:** 10.64898/2026.09.21.753290

**Authors:** Stephen J. Staklinski, Adam Siepel

## Abstract

Single-cell transcriptomics has transformed our ability to characterize cellular identity, but present-day gene expression profiles capture only a snapshot of a process that unfolds across cell division history. Recent single-cell lineage-tracing technologies make it possible to reconstruct cell division histories for thousands of cells, opening a window into how gene expression evolves. Yet observed gene expression is often redundant, with correlations among genes reflecting underlying latent biological programs and regulatory networks. To capture this structure, we introduce scPFA, a single-cell phylogenetic factor analysis framework that represents gene expression through a small set of latent factors that evolve along lineages under a phylogenetic prior, capturing correlated structure hidden in present-day observations alone. Simulations demonstrate accurate recovery of latent factors and covariance structure across conditions. Applied to developmental and cancer lineage-tracing datasets, the model uncovers biologically interpretable, lineage-associated expression programs. Together, these results demonstrate how lineage-informed factor analysis can reveal temporal biological structure hidden within high-dimensional single-cell data.

## Introduction

Gene expression is a key way in which cellular identity, differentiation, and disease progression are measurable. A cell’s transcriptional state reflects its type, its environment, its response to cell-cell interactions or signaling within that environment, and the regulatory programs active within it. Single-cell transcriptomics has made this variation visible at enormous scale [1, 2], showing that tissues and tumors are not only collections of discrete cell types, but also mixtures of continuously varying molecular states [3–6]. This heterogeneity is central to many biological processes, from development and tissue homeostasis to cancer progression and therapeutic response [7–10]. Yet a transcriptional profile remains a snapshot. It describes the state of a cell at the moment it was measured, while the processes that produced that state unfolded through cell division, differentiation, migration, and regulatory change over time [5, 11, 12].

Much of recent single-cell analysis can be viewed as an effort to recover some of this missing temporal context [13]. Trajectory inference methods arrange cells along putative developmental paths, often defined in terms of a pseudotime, by using similarities in their observed transcriptional profiles, whereas RNA velocity uses a biophysical model of local RNA splicing dynamics, based on nascent and mature transcript abundances, to infer the likely short-term direction of movement through transcriptional space [7,11,14–16]. These methods have provided powerful ways to reason about dynamic processes from static measurements, but they do not directly consider the ancestry of the cells being analyzed and they often rely on strong assumptions about the biological process being studied [12, 17, 18]. Explicit lineage-tracing efforts add a different kind of temporal information by capturing a record of cell division history through artificial barcodes, naturally occurring somatic mutations, or other markers which are claimed to be heritable [19–24]. When coupled with single-cell RNA sequencing, these technologies make it possible to observe a cell’s present-day gene expression together with a partial record of the history that produced it [12, 25].

Lineage-resolved transcriptomic data naturally invite models of gene expression on lineage trees. This follows from the covariance structure of the data, a point made early in comparative methods for phylogenetically structured data [26]. In organismal biology, stochastic processes defined along phylogenies have long been used to model the evolution of continuous quantitative traits, including gene expression, while accounting for shared evolutionary history [26–36]. Similar ideas are now being adapted to cell-lineage data to study transcriptional heritability and reconstruct changes in gene expression across time [37–40]. These approaches suggest that lineage information can reveal structure that is difficult to identify from present-day measurements alone.

One reason this hidden temporal structure may be biologically meaningful is that cellular phenotypes are rarely controlled by isolated genes acting alone. They are shaped by coordinated transcriptional programs, shared regulatory mechanisms, signaling pathways, and latent biological processes that affect many genes at once over time [41–44]. As a result, the field often assumes that the high-dimensional and noisy variation observed across thousands of genes often has lower-dimensional structure [45–48]. If these latent programs are themselves inherited, altered, or constrained along cell lineages, then treating genes independently may miss the level at which transcriptional heritability is well organized. The relevant object may be the inheritance of gene modules and the cellular states they define, not only the inheritance of individual genes [49, 50].

Single-gene models are therefore limited in their ability to represent the correlation structure among genes. One idea has been to use network-based module clustering analyses to identify heritable transcriptional signals carried by groups of genes [51]. This approach reveals gene modules, but it does not directly model how individual cells are represented by that heritable module structure. Answering that question would require post hoc integration with a separate method for learning latent representations of single cells [46] or a gene program scoring approach that assigns a score to each module in every cell [52]. Conversely, several methods learn lineage-informed embeddings or trajectories of cells, but do so without explicitly modeling the interpretable gene modules that give rise to that latent structure [53–58]. Similarly, active work has uncovered the heritability of predefined discrete cellular states along lineage trees, but without learning what gene modules those states represent within the same model [59–64]. Together, these approaches reflect substantial interest in lineage-informed gene expression structure and heritability, but each captures only one part of what appears to be a connected problem of discovering heritable gene modules while also modeling the historical representation of cells in terms of those modules. This idea is related to recent work extending RNA velocity to consider gene regulatory networks [65], but our goal is to use cell-lineage information to learn a global representation of temporal gene expression structure rather than local transcriptional dynamics.

Factor analysis provides a natural language for this problem. Originally developed in the social sciences to explain correlations among measured variables using a smaller number of unobserved factors, factor analysis provides a rich general statistical framework for learning latent low-dimensional structure in multivariate data [66]. The key to factor analysis is that it not only learns the loading matrix defining the composition of underlying latent factors, but also learns a factor matrix of those factors that provides a latent representation for each individual in the observed data. Factor analysis is not new in cell biology. In single-cell genomics, factor analysis has been adapted in various ways for deconvolving bulk gene expression data into single-cell profiles [67]. A version of factor analysis has also been recently adapted for learning cellular ecosystems from single-cell transcriptomics data [68]. In parallel, work in evolutionary biology has combined factor analysis of traits with phylogenetic models to account for shared ancestral relationships [69–71]. These efforts are part of a broader literature showing the value of learning low-dimensional structure in high-dimensional correlated traits evolving along phylogenies [72, 73]. Together, they suggest that factor analysis models are a natural fit for lineage-resolved single-cell gene expression, although, to the best of our knowledge, they have not yet been applied in this setting.

Here, we introduce scPFA (single-cell Phylogenetic Factor Analysis), a phylogenetic factor analysis framework for lineage-resolved single-cell gene expression. The model represents transcriptional variation through a smaller set of latent factors, fits stochastic processes for those factors along a cell-lineage tree, and learns the gene modules associated with each factor in a unified probabilistic framework. Using simulated and experimental lineage-tracing datasets, we work through several technical considerations and demonstrate that this approach can recover latent programs representing heritable modules of coordinated gene expression. In experimental data, the inferred factors capture biologically coherent programs that are reproducible across clones within the same system but distinct across different systems. These factors can also integrate modules identified as distinct by existing clustering methods into a single heritable axis of variation when they share similar evolutionary trajectories.

## Results

### A phylogenetic factor analysis model of heritable structure in single-cell gene expression

The goal of scPFA is to represent lineage-resolved gene expression at the level at which we expect much of the biology to be organized. We do not treat genes as independent, and we do not aim only to learn a minimally interpretable embedding of cells. Instead, we represent expression through interpretable latent transcriptional programs whose activities vary across the cell lineage. This framework is inspired by previous work on learning low-dimensional representations of single-cell gene expression [46] and on modeling the evolution of those representations along cell-lineage trees [54]. The distinction here is that the factors remain tied directly to genes, so that each latent axis can be interpreted as both a gene module and a heritable cellular state variable (**Fig. 1A**).

**Figure 1:**
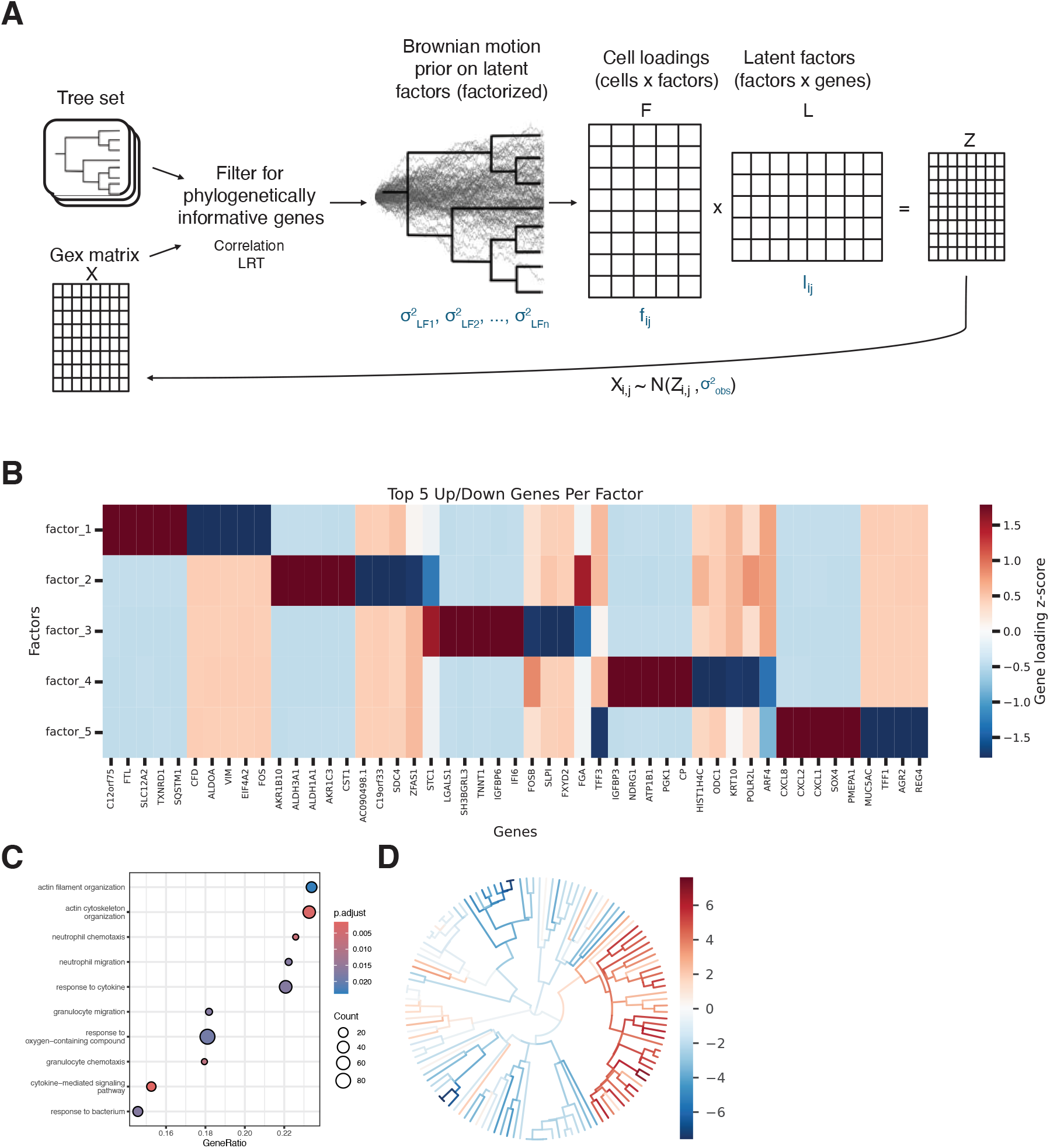
Overview of scPFA and representative visualizations. **(A)** The scPFA model. Genes are first preprocessed to identify those with phylogenetic signal, and then latent factors are modeled on the tree using independent Brownian motion priors, yielding a cell-factor matrix **F** and factor-gene loading matrix **L**, whose product reconstructs gene expression. Blue variables denote model parameters inferred during fitting, whereas black variables are fixed. **(B)** Example factor-gene loading matrix **L**, shown only for the highest positive and negative gene loadings for each factor. **(C)** Gene set enrichment analysis of genes associated with one inferred factor, illustrating biological interpretation of latent gene programs. **(D)** Reconstruction of a latent factor across the lineage tree using the inferred cell-factor matrix **F**, enabling visualization of temporal dynamics along cellular histories.

As a first formulation, we work with a log-transformed matrix of normalized expression values, **X** ∈ ℝ^*n*×*p*^, for *n* cells and *p* genes, together with a cell-lineage tree *T*. We treat this matrix as a continuous trait matrix under a Gaussian observation model, and restrict the analysis to genes with evidence of phylogenetic signal. This does not model the unique observational properties of single-cell count data, as we have done in related prior work [40], but instead focuses the problem more generally on whether utilizing lineage information in a factor analysis framework helps recover interpretable low-dimensional structure in gene expression.

Observed expression is modeled as a noisy low-rank reconstruction, with **X** = **Z** + **E** and **Z** = **FL**. Here **F** ∈ ℝ^*n*×*k*^ is a cell-by-factor matrix, **L** ∈ ℝ^*k*×*p*^ is a factor-by-gene loading matrix, and **E** represents residual noise, with 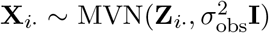. The rows of **L** define gene modules, while the columns of **F** describe the activity of those modules across cells.

The lineage enters through a Brownian motion prior on the columns of **F**. For each factor *d*, we place an 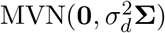 prior on **F**_*d*_, where **Σ** is the covariance matrix induced by the lineage tree and 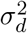 is a factor-specific Brownian variance parameter learned from the data. This prior makes factor activities more similar among cells with more shared history, while allowing divergence to accumulate along longer or independent branches. The variance parameters determine the extent to which each latent factor varies across the lineage, allowing different transcriptional programs to evolve at different rates. More complex evolutionary models could be used, but Brownian motion provides a simple and interpretable baseline, avoids additional process parameters, and is consistent with cautionary work showing that parameter-rich alternatives such as Ornstein–Uhlenbeck models do not necessarily improve inference in practice [74].

The inferred quantities are **F, L**, the factor-specific Brownian variance parameters 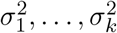, and 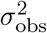 (blue variables in **Fig. 1A**). The main biological output is **L**, which defines gene modules associated with each latent factor (**Fig. 1B**). Factors can be annotated by ranking genes according to their positive and negative loadings and performing gene set enrichment analysis (**Fig. 1C**). The inferred matrix **F** can also be used to reconstruct each latent program across cellular history (**Fig. 1D**). As in any factor analysis model, the factorization has scale, sign, and permutation invariances. In the empirical work below on simulated data, we introduce the use of constrained **L** rows and post hoc rules as sufficient means for resolving these invariances.

### Identifying phylogenetically informative genes prior to factor analysis

Matched lineage-tracing and transcriptomic data make it possible to ask which genes vary in a way that is consistent with shared lineage history. Because our model is intended to learn heritable gene modules, we first filtered to genes with evidence of phylogenetic signal. This is a simple approximation to a more general model in which heritable and non-heritable components would be learned jointly.

We compared PATH, a recently proposed correlation-based filter [61], with two likelihood-ratio tests based on Brownian motion covariance models. The first LRT used an empirically estimated null distribution, whereas the second used a computationally efficient Pagel’s *λ* parameterization [75, 76] like that which we have used in prior work [40]. We also re-implemented PATH in C for convenience within our workflow, which gave numerically identical results to the original R implementation but ran faster.

In simulations with known heritable genes, PATH was precise but less sensitive than either LRT (**Fig. 2A**). The LRTs recovered nearly all simulated heritable genes, as expected under data generated from Brownian motion, but differed in runtime. The empirical-null LRT became expensive as dataset size increased, whereas the *λ*-LRT gave similar accuracy at much lower cost due to the analytical null distribution. We therefore find that it is best to use the *λ*-LRT for datasets up to approximately 1,000 cells and view PATH as a practical alternative for larger datasets, with the caveat that it may discard some informative genes.

**Figure 2:**
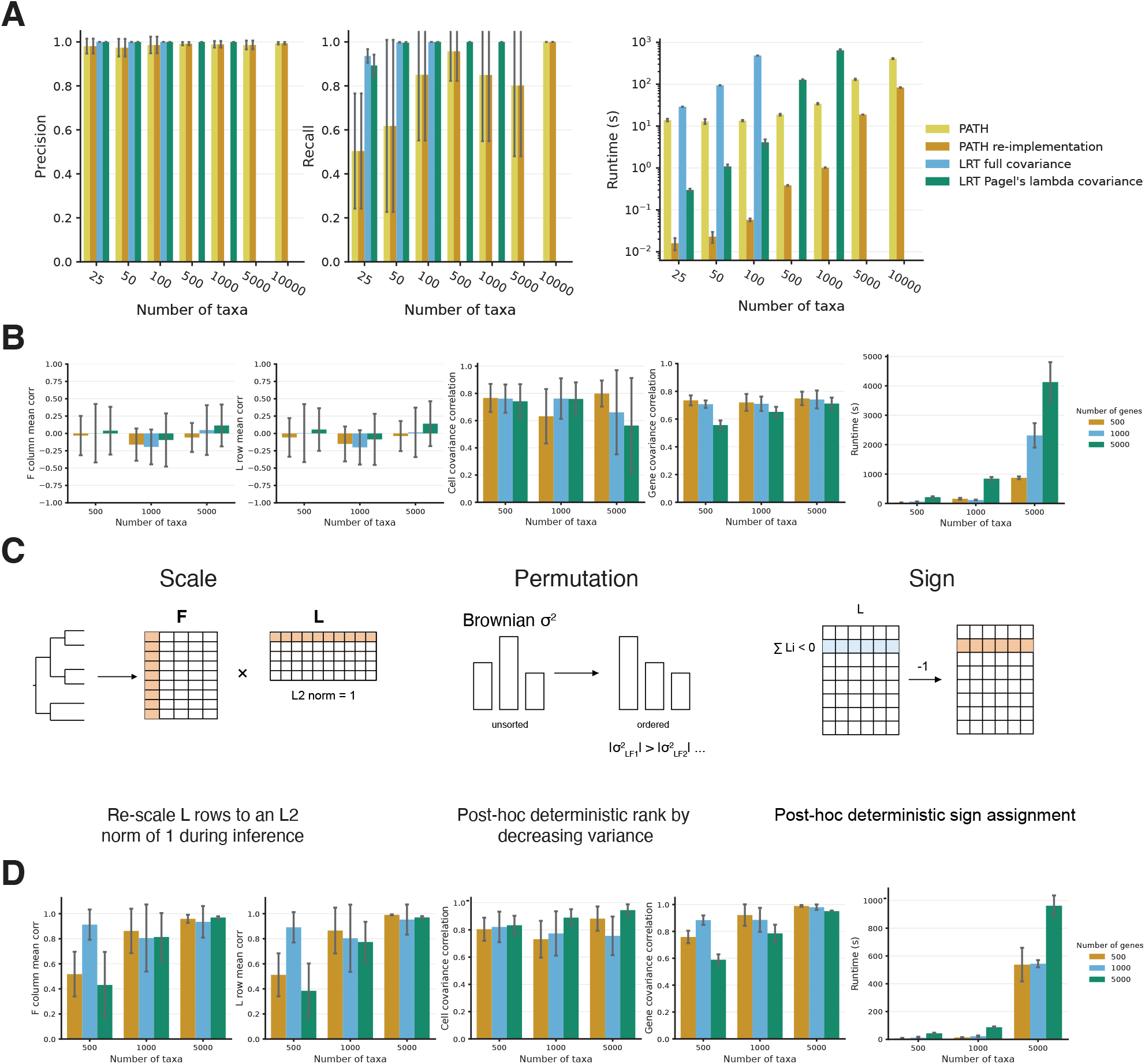
Simulation benchmarking and identifiability constraints for phylogenetic factor analysis. **(A)** Performance, in terms of precision (left), recall (middle), and runtime (right), of four methods for identifying phylogenetically informative genes across simulated datasets. Methods include the original PATH correlation-based approach, a reimplementation of PATH, and two likelihood-ratio test (LRT) approaches based on Brownian motion covariance models. **(B)** Recovery of latent factors before resolving model invariances. From left to right: mean Pearson correlation between inferred and true columns of **F**, mean Pearson correlation between inferred and true rows of **L**, cell-cell covariance correlation, gene-gene covariance correlation, and runtime across dataset sizes. **(C)** Graphical representation of constraints used to resolve scale, permutation, and sign invariances in the factorization. **(D)** Model performance after applying invariance constraints and refitting. Panels are shown in the same order as in (B), demonstrating improved recovery and interpretability of latent factors while maintaining accurate covariance reconstruction.

PATH’s reduced recall motivated us to ask whether its performance depended on properties of the simulated lineage trees. The strongest pattern was for trees with very short branches, where nearby cells would be expected to have weak Brownian divergence relative to observational noise (**Supplementary Fig. S1**). Nonetheless, the impact on recall was modest, particularly on larger trees, and PATH’s precision remained high.

### Constraining factor analysis invariances reproducibly recovers simulated latent structure

Factor analysis models are inherently non-identifiable. The same low-rank reconstruction can be written after rescaling, permuting, or changing the signs of the factors amongst other rotational invariances. In simulations, before resolving these invariances, individual columns of **F** and rows of **L** were not recovered consistently, even though the induced cell-cell and gene-gene covariance structures were recovered reasonably well (**Fig. 2B**). Post hoc alignment to the generating factors largely corrected this problem, confirming that much of the error reflected factor-basis ambiguity rather than failure to recover the latent structure (**Supplementary Fig. S2A**).

We therefore added simple constraints to make the fitted factors more reproducible (**Fig. 2C**). Rows of **L** were constrained to have unit *ℓ*_2_ norm by rescaling them during inference, factors were ordered by decreasing Brownian variance post-inference, and signs were chosen deterministically from the sum of factor loadings post-inference. These choices were aimed at fixing scale, permutation, and sign without changing the model’s basic structure. An alternative would be to fix the Brownian variance parameters to 1, as in prior phylogenetic factor analysis work [70, 71]. In our setting, however, this gave poorer recovery than normalizing the rows of **L**. This difference may reflect our maximum a posteriori optimization, where the factor-gene parameters are learned by stochastic gradient descent. Fixing the Brownian variances only changes the effective prior on **F**, whereas row normalization provides a more direct constraint on the fitted loading matrix **L**.

With these constraints in place, the fitted factors more closely matched the generating **F** and **L** across simulated dataset sizes, with moderate additional gains in recovery of the cell-cell and gene-gene covariance structure (**Fig. 2D**). It is also notable that the constraints seemed to substantially improve runtime. Post hoc alignment seemingly also had little left to improve, consistent with the main factorization ambiguities having been removed (**Supplementary Fig. S2B**). Representative simulations showed the same behavior at the level of the simulated **F, L, Z**, and **X** matrices (**Supplementary Fig. S3A**) and in low-dimensional views of the latent and observed expression states (**Supplementary Fig. S3B**).

Recovery was generally stable across simulated dataset sizes, with the main exception occurring for the smallest trees paired with the largest gene sets. In the 500-cell, 5,000-gene setting, recovery was weaker (**Fig. 2D**), a pattern consistent with prior work showing that inference becomes more difficult when the number of traits exceeds the number of observations [73]. Interestingly, we did not observe catastrophic failure in this regime, just minor underperformance. We do not pursue this limitation further here because lineage-tracing datasets are moving toward larger trees, whereas the number of genes expressed by any cell is biologically constrained. Thus, the natural data regime for this problem is one with many more cells than genes, where the model seemingly performs well.

### Regularization encourages distinct gene modules in real data

We next aimed to apply our model to two lineage-tracing datasets, one in a developmental biology setting [62] and the other in lung cancer [10]. Based on the simulation results above, we restricted these analyses to clones containing more than 1,000 cells, where reconstruction was generally strongest. This filtering left two clones from the developmental dataset and four clones from the lung cancer dataset.

We initially fit the model using the invariance constraints established in simulations. Although these constraints encouraged the factorization to be reproducible, the resulting factors often contained overlapping gene loadings that were difficult to interpret (**Fig. 3A**). In many respects, the goal of our model is not simply to reconstruct expression from a low-rank representation, but to identify relatively distinct and interpretable factors corresponding to biologically meaningful combinations of genes. We therefore sought to constrain two potential sources of factor overlap: insufficient sparsity within rows of **L** and similarity in the directions that different rows of **L** point through “gene space”.

**Figure 3:**
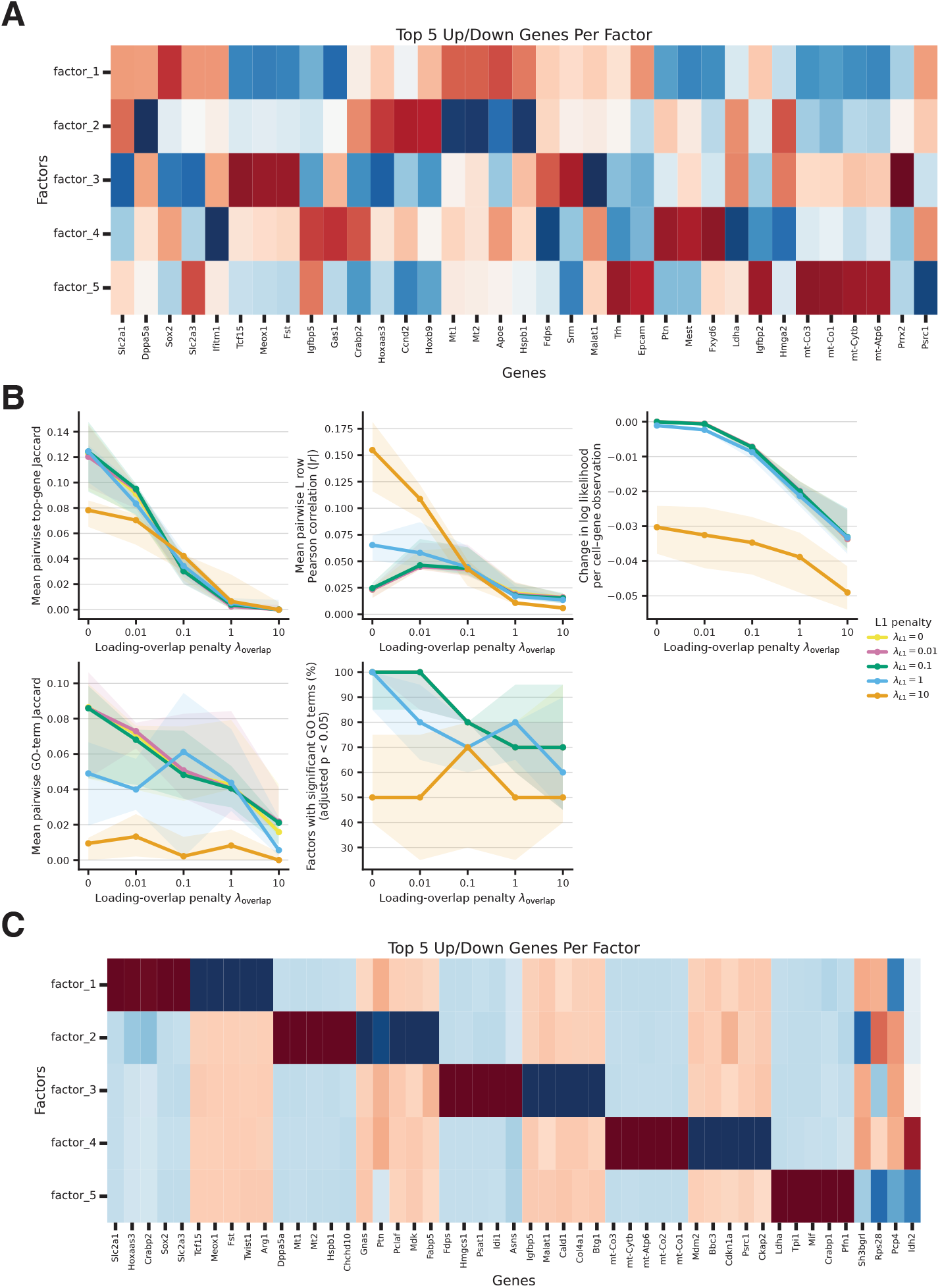
Regularization encourages distinct factors in real data. **(A)** Factor-gene loading matrix **L** before constraining factor similarity, showing the five largest positive and negative gene loadings per factor. **(B)** Effects of varying row-wise *ℓ*_1_ and cosine-similarity overlap penalties. Lines and shaded regions show the mean and interquartile range across datasets, respectively. Metrics are pairwise Jaccard similarity of the top 10 positive and negative genes per factor, ignoring sign (top left); pairwise Pearson correlation between rows of **L** (top middle); dataset-size-normalized log-likelihood (top right); pairwise Jaccard similarity of Gene Ontology (GO) terms (bottom left); and the percentage of factors with at least one significant GO term (bottom middle). **(C)** Loading matrix after applying the selected penalties of *ℓ*_1_ *λ*_1_ = 0.1 and cosine-similarity overlap *λ*_*C*_ = 1, displayed as in (A).

To address these sources of overlap, we implemented dataset-size-adjusted penalties consisting of an *ℓ*_1_ penalty on each row of **L** and a cosine-similarity overlap penalty between rows. We evaluated their effects on numerical factor overlap using top-gene Jaccard similarity and row-wise Pearson correlation, on model fit using normalized log-likelihood, and on biological function overlap using shared Gene Ontology (GO) terms and the fraction of factors with significant GO enrichment (**Fig. 3B**).

The *ℓ*_1_ penalty had little consistent effect on either numerical or biological overlap, other than substantially reducing model fit at high values. In contrast, the overlap penalty meaningfully reduced both top-gene overlap and correlation between rows of **L**, while also tending to reduce overlap between their associated GO terms. We therefore selected an overlap penalty of *λ*_*C*_ = 1 and an *ℓ*_1_ penalty of *λ*_1_ = 0.1. The modest *ℓ*_1_ penalty was retained to encourage some sparsity, although its empirical contribution appeared limited and we recognize that this may be partly redundant. Under these conditions, the fitted loading matrices contained substantially more distinct and interpretable factors (**Fig. 3C**).

For these tests, we fixed the number of factors to five across all six clones. However, the percentage of factors with significant GO enrichment was often below 100% (**Fig. 3B**), and the last factor in the representative loading matrix appeared to have relatively weak structure (**Fig. 3C**). These observations suggested that five factors may not be appropriate for every clone. Choosing dimensionality is a common challenge in similar tasks, and these results motivated us to consider it more carefully in the final model fits for each dataset.

### Lineage-informed factors reveal conserved developmental transitions and clone-specific cell-cycle programs

The developmental lineage-tracing dataset was generated using a mouse embryonic stem cell model of the trunk that proceeded through four days of gastrulation followed by one day of neural tube differentiation [62]. We selected the two largest clones, trunk-like structures 1 and 2 (TLS1 and TLS2), each containing more than 1,000 cells. We first applied the correlation-based PATH filter to identify genes with phylogenetic signal. A large fraction of ribosomal and mitochondrial genes passed this filter (**Supplementary Fig. S4A**), consistent with prior work challenging the assumption that ribosomal genes are static housekeeping genes [50]. Retaining these genes and fitting the factor analysis model produced a single dominant ribosomal and mitochondrial factor that obscured structure in other modules, so we removed them from subsequent analyses.

Among the remaining genes, 8.4% in TLS1 and 15.8% in TLS2 showed phylogenetic signal (**Supplementary Fig. S4B**). Of the genes shared between samples, 5.8% showed signal in both, with an additional 3% specific to TLS1 and 11% specific to TLS2. Despite this limited gene-level overlap, the two sets were enriched for nearly identical GO biological processes, including pattern specification, embryonic organ development, morphogenesis, and stem cell differentiation (**Supplementary Fig. S4C**). Thus, different genes carried phylogenetic signal in the two clones, but they converged on the same developmental processes.

We next fit the phylogenetic factor analysis model to the filtered genes from each clone. To select the latent dimensionality *k*, we evaluated log-likelihood, significant GO enrichments, newly identified GO terms, and the number of factors explaining most of the variance under the Brownian motion prior across a range of values. Improvements in both clones plateaued near *k* = 10 (**Supplementary Fig. S5**). At larger values of *k*, the model collapsed excess factors by shrinking their Brownian variance to the specified floor. We therefore selected *k* = 10 for both clones.

Only four factors in TLS1 and six in TLS2 had Brownian variance above the floor in the final model fits (**Fig. 4A,C**). The factors had distinct top genes (**Fig. 4B,D**) and direction-specific GSEA enrichments within each clone (**Supplementary Fig. S6A**). Comparison of their gene loadings across clones identified three conserved programs (**Fig. 4E**). The strongest correspondence was between TLS1 factor 4 and TLS2 factor 2 (*r* = 0.93). In both clones, one pole contained *Dppa5a, Chchd10*, and *Ckb*, whereas the opposing pole contained *Mdk, Gnas, Ldha, Fabp5*, and *Hmga2*. TLS1 factor 4 was positively enriched for leukemia inhibitory factor (LIF) response and negatively enriched for gliogenesis and spinal cord development. These results suggest a reproducible lineage-varying axis between a stem-cell-associated state and neural differentiation.

**Figure 4:**
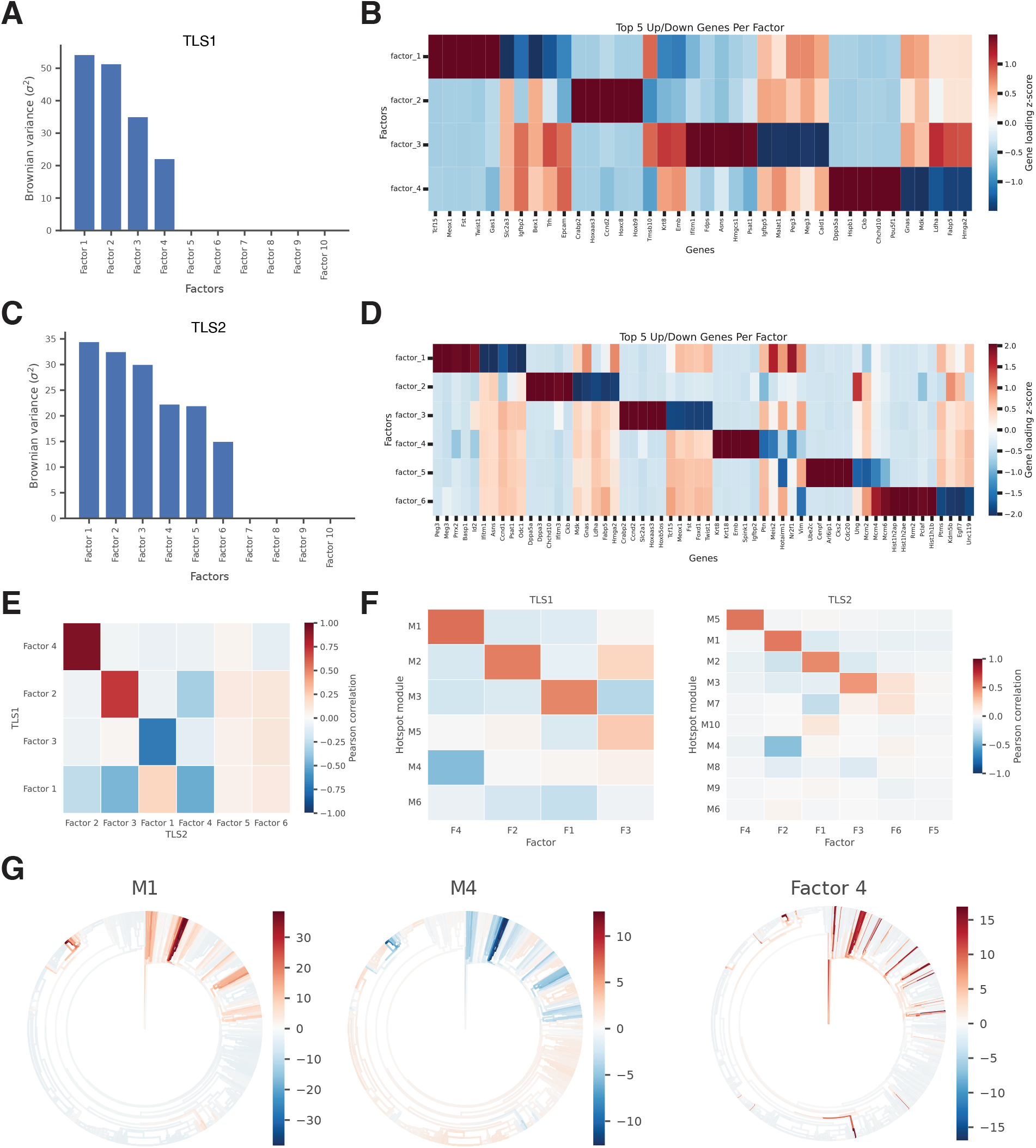
Phylogenetic factors recover developmental gene expression programs. **(A)** Fitted Brownian variance (*σ*^2^) across 10 TLS1 factors. **(B)** Standardized loadings for the five largest positive and negative genes per active TLS1 factor. **(C–D)** Brownian variances and gene loadings for TLS2, displayed as in (A–B). **(E)** Pearson correlations between active TLS1 and TLS2 factor-loading profiles. **(F)** Pearson correlations between factor-loading profiles and Hotspot modules in TLS1 (left) and TLS2 (right). **(G)** Representative collapse of TLS1 Hotspot modules M1 and M4 into factor 4, with module and factor scores projected along the lineage tree.

TLS1 factor 2 aligned with TLS2 factor 3 (*r* = 0.71). The corresponding poles in both factors contained *Crabp2, Ccnd2, Hoxaas3*, and posterior *Hox* genes, including *Hoxc8, Hoxb9*, and *Hoxb5os*. In TLS1 factor 2, this pole opposed epithelial-associated genes including *Epcam, Igfbp2, Krt8*, and *Emb*. In TLS2 factor 3, it opposed *Tcf15, Meox1, Fst, Foxd1*, and *Twist1*. Thus, the same posterior-patterning program was recovered in both clones but was coupled to different alternative transcriptional states.

TLS1 factor 3 corresponded inversely to TLS2 factor 1 (*r* = −0.71), reflecting an arbitrary reversal of factor orientation. In both cases, one pole was marked by *Ifitm1, Asns*, and *Psat1*, whereas the other contained *Peg3, Meg3, Prrx2*, and *Basp1*. The associated enrichments separated small-molecule and organophosphate metabolism from organ and tube morphogenesis, adhesion, motility, and vascular development. TLS1 factor 3 and TLS2 factor 1 therefore recovered the same coupling between biosynthetic activity and morphogenetic state despite their opposite signs.

The remaining active factors captured clone-specific variation. TLS1 factor 1 was marked on one side by *Tcf15, Meox1, Fst, Twist1*, and *Gas1* and on the other by *Slc2a3, Igfbp2, Bex1, Trh*, and *Epcam*. Its enrichments linked the first pole to pattern specification and muscle development and the second to carbohydrate transport. In TLS2, factor 4 was distinguished by the epithelial genes *Krt8, Krt18, Emb, Spink1*, and *Igfbp2*. TLS2 factors 5 and 6 instead resolved proliferation into two programs. Factor 5 contained mitotic genes including *Ube2c, Cenpf, Arl6ip1, Cks2*, and *Cdc20* and was strongly enriched for cell division and organelle fission. Factor 6 was marked by replication-associated histones together with *Rrm2* and *Pclaf* and was enriched for DNA replication, repair, and chromosome organization. The TLS2-specific signal therefore included an epithelial program and separately heritable mitotic and S-phase programs rather than a single generic cell-cycle axis.

Finally, we compared the fitted factors with the mutually exclusive gene modules that can be recovered through network-based clustering using a method named Hotspot [51]. Hotspot constructs a K-nearest-neighbor similarity graph between cells, in this case based on lineage distance, and identifies gene modules by clustering genes with locally correlated expression patterns across neighboring cells in the graph. Several factors had direct Hotspot counterparts (**Fig. 4F**). TLS1 factors 4, 2, and 1 aligned most strongly with Hotspot modules M1, M2, and M3, respectively, while TLS2 factors 4, 2, 1, and 3 aligned with M5, M1, M2, and M3. However, individual factors could also collapse multiple Hotspot modules. For example, TLS1 factor 4 was positively associated with M1 and negatively associated with M4, placing two mutually exclusive modules on opposing sides of the conserved stem-cell-to-differentiation axis. These modules exhibited coordinated changes along the lineage (**Fig. 4G**), supporting their representation as components of a single lineage-varying program. Thus, Hotspot recovered many of the same local gene associations, while the factor model organized them into signed developmental axes and separately identified the TLS2 mitotic and DNA-replication programs.

### Lineage-informed factors reveal shared and clone-specific programs in lung cancer progression

The lung cancer dataset of interest contained four clones with more than 1,000 cells, ranging from ∼1,000 cells in CP4 to ∼10,000 cells in CP1, resulting from a KRAS-mutant lung adenocarcinoma cell line (A549) xenograft model that progressed to metastasis over 54 days [10]. As in the developmental dataset, ribosomal and mitochondrial genes frequently showed phylogenetic signal (**Supplementary Fig. S7A**) and contributed strongly to a single dominant factor when retained in downstream factor analysis. We therefore removed these genes before subsequent analyses.

Among the remaining genes, the fraction with phylogenetic signal varied from 1.4% in CP3 to 9.2% in CP1 (**Supplementary Fig. S7B**). Of the 16,773 genes detected in all four clones, 2,075 showed phylogenetic signal in only one clone, whereas only 26 showed phylogenetic signal in all four. The associated GO enrichments also differed across clones. CP1 was dominated by oxidative phosphorylation, wound healing, and cell migration; CP2 by apoptosis, reactive oxygen species metabolism, and detoxification; and CP4 by immune responses and detoxification. CP3 had no significant GO enrichment among its smaller filtered gene set (**Supplementary Fig. S7C**). Thus, the cancer clones shared some broad stress-response processes but differed substantially in both the number and identity of genes carrying phylogenetic signal.

We next fit models across a range of latent dimensions using the same selection criteria as for the developmental data (**Supplementary Fig. S8**). CP1 and CP2 were the two largest clones and retained the most genes with phylogenetic signal, whereas CP3 retained few genes, providing limited support for fitting a multifactor model. We therefore focused subsequent factor analysis on CP1 and CP2. For both clones, Brownian variance was concentrated in relatively few factors, and excess factors were commonly shrunk to the variance floor. We selected *k* = 10 as a common upper bound. Four factors in each clone retained Brownian variance above the floor in the final model fits (**Fig. 5A,C**). These factors had distinct top genes (**Fig. 5B,D**) and direction-specific GSEA enrichments within each clone (**Supplementary Fig. S9**).

**Figure 5:**
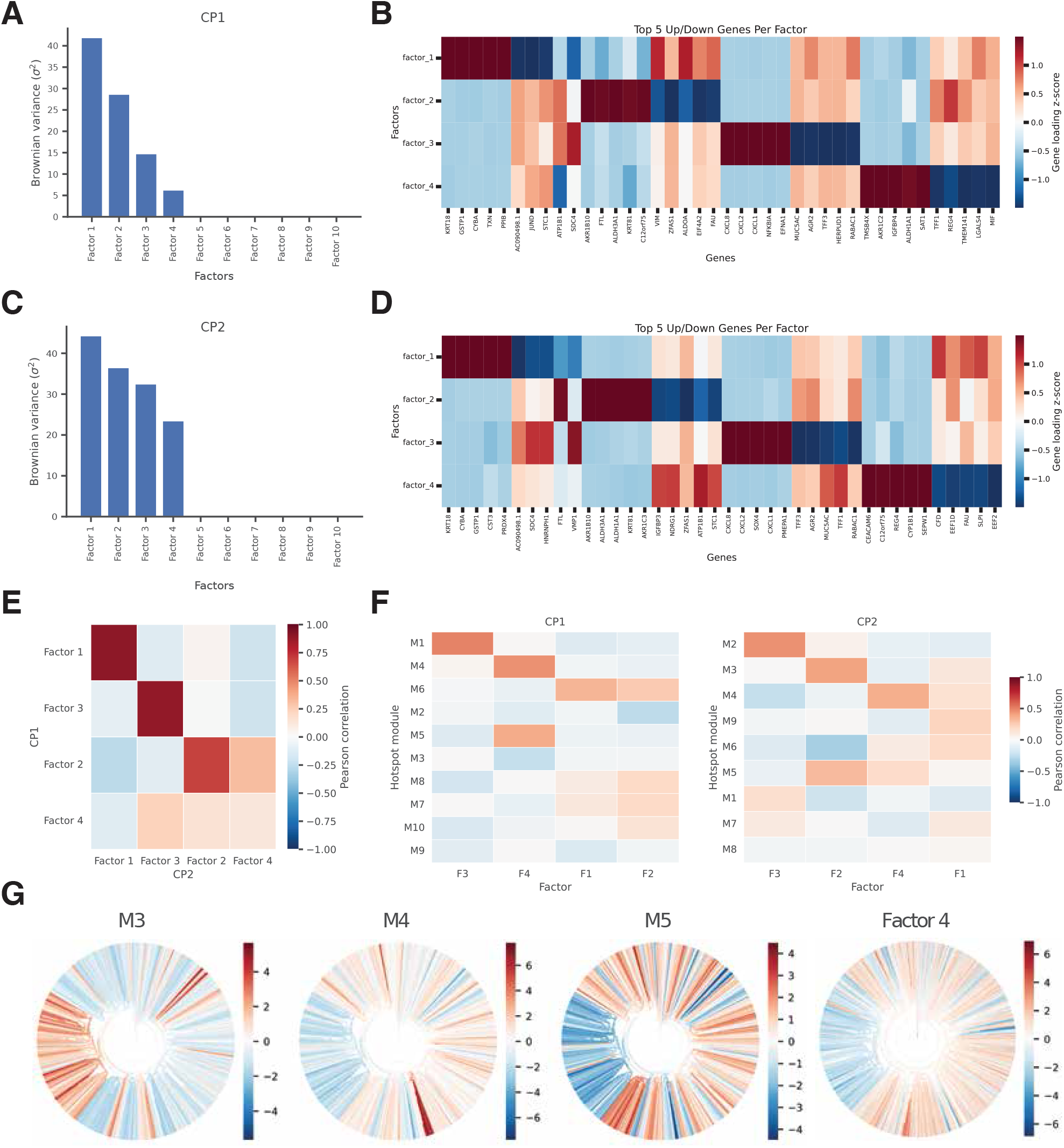
Phylogenetic factors recover gene expression programs in lung cancer. **(A)** Fitted Brownian variance (*σ*^2^) across 10 CP1 factors. **(B)** Standardized loadings for the five largest positive and negative genes per active CP1 factor. **(C–D)** Brownian variances and gene loadings for CP2, displayed as in (A–B). **(E)** Pearson correlations between active CP1 and CP2 factor-loading profiles. **(F)** Pearson correlations between factor-loading profiles and Hotspot modules in CP1 (left) and CP2 (right). **(G)** Representative collapse of CP1 Hotspot modules M3, M4, and M5 into factor 4, with module and factor scores projected along the lineage tree.

Three of the four active factors in CP1 had clear counterparts in CP2 (**Fig. 5E**). The strongest correspondence connected CP1 factor 1 with CP2 factor 1 (*r* = 0.89). Both factors had a pole marked by *KRT18, GSTP1, CYBA*, and *PRDX4*, consistent with an epithelial oxidative-stress program. CP1 factor 3 aligned with CP2 factor 3 (*r* = 0.88) and was marked in both clones by *CXCL8, CXCL2, CXCL1, NFKBIA*, and *PMEPA1*. The opposing poles contained secretory epithelial genes including *MUC5AC, AGR2*, and *TFF3*. Thus, both clones contained a closely matched lineage-varying program separating an inflammatory chemokine state from a secretory epithelial state. A third correspondence linked CP1 factor 2 and CP2 factor 2 (*r* = 0.68). Both were distinguished by *AKR1B10, ALDH3A1, KRT81*, and related detoxification genes. In contrast, CP1 and CP2 factors 4 showed little similarity (*r* = 0.11). CP1 factor 4 separated *AKR1C2, IGFBP4, ALDH1A1*, and *SAT1* from *TFF1, REG4*, and *LGALS4* and was enriched for morphogenesis and organ development. CP2 factor 4 instead had a pole marked by *CEACAM6, C12orf75, REG4*, and *CYP1B1* and was enriched more broadly for signaling and cell communication. The two largest clones therefore shared inflammatory, oxidative-stress, and detoxification programs but differed in an additional lineage-varying program.

The fitted factors were again concordant with several Hotspot modules in the lung cancer dataset (**Fig. 5F**). CP1 factor 3 and CP2 factor 3 aligned most strongly with Hotspot modules M1 and M2, respectively, while the detoxification-associated factor 2 aligned with CP2 modules M3 and M5. As in the developmental data, individual factors also connected multiple mutually exclusive Hotspot modules. CP1 factor 4 captured coordinated variation represented separately by modules M3, M4, and M5, whose scores changed together along the lineage tree (**Fig. 5G**). These results support the recovered gene-level programs while showing how the signed factors organize multiple co-expression modules into shared and clone-specific axes of lineage-associated variation.

### Lineage-associated programs are distinct across developmental and lung cancer model systems

We next asked whether the factors learned in the developmental and lung cancer datasets represented similar biological processes. Because the two datasets were generated using cells from different species and retained different phylogenetically informative genes, we compared each active developmental factor with each active cancer factor using the Jaccard similarity of their significantly enriched GO biological process terms (**Fig. 6**). Most cross-context comparisons showed no overlap, with 61 of 80 factor pairs having a Jaccard similarity of zero, and even the strongest similarity was only 0.18. Thus, most learned lineage-associated factors appear to be specific to the biological system in which they were inferred, rather than reflecting a small set of generic heritable expression programs.

**Figure 6:**
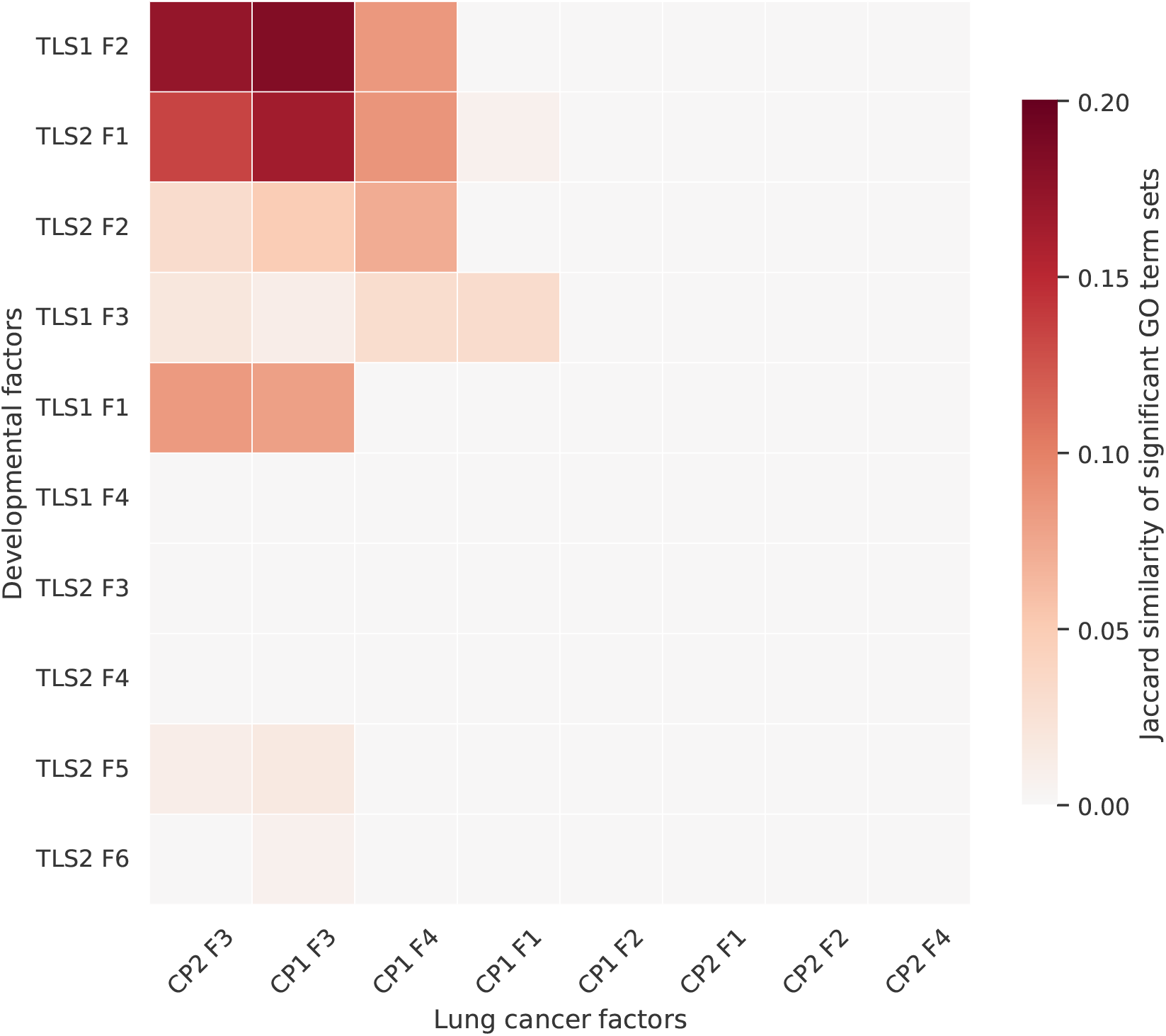
Phylogenetic factors are functionally distinct across developmental and cancer systems. Jaccard similarity between sets of significantly enriched Gene Ontology biological process terms for each active developmental factor (rows) and active lung cancer factor (columns). Factors are ordered around the strongest one-to-one matches.

The few detectable similarities largely involved broad processes that are known to be engaged in both development and cancer. TLS1 factor 2 overlapped most strongly with CP1 and CP2 factor 3 (Jaccard similarities of 0.18 and 0.17, respectively), sharing terms related to transcriptional and biosynthetic regulation, cell differentiation, neurogenesis, and organ development. TLS2 factor 1 also overlapped with CP1 and CP2 factor 3 (0.16 and 0.13), with shared processes including cell migration and adhesion, cytoskeletal organization, wound response, cell differentiation, morphogenesis, and vascular development. Weaker correspondences linked developmental factors to cancer programs through neural, epithelial, tube, and circulatory-system morphogenesis, while TLS2 factors 5 and 6 shared only general cell-cycle terms with cancer factor 3. These limited overlaps suggest that the common signal is concentrated in general regulatory, proliferative, and morphogenetic processes, whereas the more specific composition of most lineage-varying programs remains context dependent.

An analogous comparison of GO enrichments for Hotspot modules gave a similar result. Of the 240 developmental–cancer module pairs, 141 shared no significant terms, and the strongest Jaccard similarity was only 0.13 (**Supplementary Fig. S10**). Thus, the cross-system functional distinctness was also evident among modules learned by an independent lineage-aware network-based clustering method.

## Discussion

We introduced scPFA, a phylogenetic factor analysis model that jointly learns lineage-varying representations of cells and the gene modules that define them. Results from fitting the model to simulated data showed that it could recover latent factors and their implied covariance structure, while additional regularization produced more distinct and interpretable factors in real data. Applied to developmental and lung cancer lineage-tracing datasets, the model identified a small number of factors with substantial Brownian variance, biologically coherent gene loadings, and significant functional enrichments that aligned well with expectations for each respective model system. Several factors were reproducible across clones and aligned with modules identified by Hotspot, while others captured clone-specific programs or combined multiple Hotspot modules into a single signed axis of lineage-associated variation.

Our gene-filtering analyses identified a small percentage of genes with detectable phylogenetic signal across both model systems, consistent with evidence for heritable transcriptional memory in single cells [49]. However, most genes showed no such signal, suggesting that gene-level heritability across the entire lineage tree is not pervasive under our chosen gene-filtering model and significance thresholds. Ribosomal and mitochondrial genes were notable exceptions, frequently exhibiting strong phylogenetic signal. This result agrees with recent evidence that many broadly expressed genes, including ribosomal genes that are often considered to be housekeeping genes, may retain notable lineage-dependent expression variation [50]. Interestingly, the lung cancer clones differed more strongly in the number and identity of genes with phylogenetic signal, consistent with a more constrained set of transitions in the directed developmental system and a broader range of heritable transcriptional outcomes in cancer. It is worth noting, however, that the present data do not distinguish cellular plasticity from differences in clone history, environment, or sampling, so further work will be required.

The biological composition of our learned heritable factors reflected the different settings represented by the two datasets well. In the developmental system, three programs were conserved across clones, including axes associated with stem-cell maintenance and neural differentiation, posterior patterning, and the coupling of morphogenesis to biosynthetic state. The lung cancer clones shared several programs centered on oxidative stress, inflammatory chemokine expression, secretory epithelial state, and detoxification.

Direct comparison across the two biological settings further showed that most inferred programs were context specific. More than three quarters of developmental–cancer factor pairs shared no significantly enriched GO terms, and the strongest overlaps remained modest. The exceptions involved broad transcriptional and biosynthetic regulation, differentiation, morphogenesis, migration and adhesion, vascular development, and cell-cycle control. Thus, a small core of general cellular programs may recur across systems, but most differ, or perhaps are reorganized into different heritable modules, between embryonic development and lung cancer progression.

More broadly, these results provide a proof of concept for using an explicit evolutionary model to discover heritable gene modules rather than testing genes independently or defining modules through clustering in a way that is constrained by the lineage. This formulation connects the composition of a program, its activity in individual cells, and its variation through cellular history within one joint probabilistic framework. It also creates a basis for formal hypothesis testing through comparisons of nested models. For example, Brownian motion could be compared with Ornstein–Uhlenbeck models of stabilizing regulation, models with shifts in optima or rates along particular branches, or models in which different factors respond to different lineage or environmental regimes. Related model comparisons have been useful for testing gene-expression dynamics in our prior work [40]. Extending them to latent factors would allow such hypotheses to be tested at the level of coordinated transcriptional programs. The Brownian model used here therefore serves as a simple baseline upon which richer biological process models can be built.

It is also worth noting that the inferred factors imply a lower-dimensional gene-expression space in which movement along cellular lineages is naturally reconstructed, yielding lineage-aware analogs of trajectory inference or Waddington landscapes that we did not fully explore here [77]. Here, we focused our efforts on recovering the gene modules themselves. However, the fitted factor analysis model provides much richer information and could be immediately used for a variety of other tasks including the reconstruction of these differentation landscapes and the study of discrete versus continuous dynamics of cell-state transitions.

The current implementation also has several limitations that motivate future work. First, it treats normalized log expression as a continuous Gaussian trait rather than modeling single-cell counts and their mean– variance relationship directly. Second, filtering genes before factor analysis helps computational tractability but conditions the model on a fixed set of genes with detectable phylogenetic signal. A joint model of heritable and non-heritable components could propagate uncertainty in this classification and recover weak signals that are apparent only at the module level. The present continuous model also does not naturally represent genes with binary or strongly zero-inflated expression, which could be incorporated through mixed observation models. Finally, the lineage tree was treated as known in the analyses presented here. Our code is set up to accept a collection of candidate trees and marginalize the Brownian factor priors across them, but we used a single tree per clone to keep this initial analysis simple. Evaluating that functionality on posterior samples of lineage trees will be important because our prior work has confirmed that uncertainty in reconstructed histories can meaningfully affect downstream biological conclusions [78]. Such extensions could propagate tree, expression, and module uncertainty together, although fully Bayesian phylogenetic factorization may require additional approximations to remain scalable to large single-cell datasets [70, 71]. These are all relatively direct extensions of the current work because they retain a fixed factor-loading matrix across the lineage. A more ambitious model could allow module composition itself to evolve as genes enter, leave, or change their contributions to a program. This could reveal how regulatory programs are reorganized during development and disease progression.

Together, our results demonstrate the promise of phylogenetic factor analysis for extracting interpretable, lineage-associated structure from high-dimensional single-cell data. By connecting gene modules, cellular states, and their shared histories in a unified framework, this approach provides both a practical tool for studying transcriptional heritability and a flexible foundation for increasingly realistic models of cellular evolution.

## Methods

### Phylogenetic factor model specification

Tree tips and expression profiles were restricted to their shared cells, and each tree was rescaled to unit height so that Brownian variance parameters are fit on a similar scale across datasets. Before phylogenetic gene filtering and factor-model fitting, counts were normalized to a total of 10,000 per cell to control for library size, transformed as log(1 + *x*) to stabilize their variance and make a Gaussian working approximation more reasonable, and centered across cells for each gene to allow modeling Brownian motion around a mean of 0.

For *n* cells, *p* retained genes, and *k* latent factors, we modeled the resulting expression matrix **X** ∈ ℝ^*n*×*p*^ as

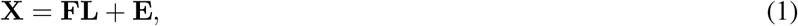

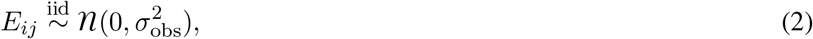

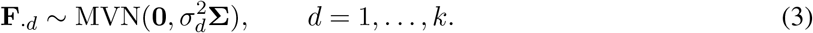

Here, **F** ∈ ℝ^*n*×*k*^ contains factor activities across cells, **L** ∈ ℝ^*k*×*p*^ contains their gene loadings, and **Σ** is the Brownian covariance induced by the lineage tree, with Σ_*ij*_ equal to the time shared by cells *i* and *j* from the root to their most recent common ancestor. Each factor has its own Brownian variance 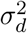, whereas 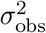 describes residual expression variation shared across genes and cells. We treated the factors as independent Brownian processes and did not estimate covariance between them, providing a simpler baseline in which lineage dependence is modeled within, but not between, factors.

Inference balances reconstructing the observed expression matrix with **FL** and requiring each column of **F** to vary across cells in a manner consistent with Brownian motion on the lineage tree. The observation variance controls unexplained expression noise, while each Brownian variance controls the amount of lineage-associated variation assigned to its factor. The observation log-likelihood is 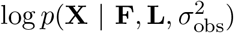. The Brownian log prior is 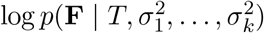. Their sum is the unpenalized joint log density used for maximum a posteriori inference of the latent factor activities:

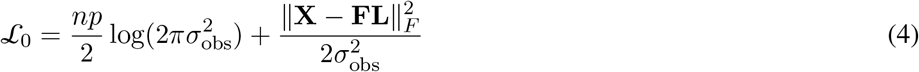

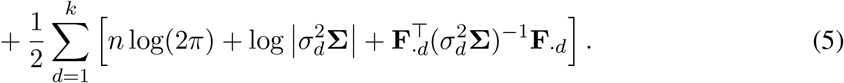

We minimized the penalized objective *L* = *L*_0_ + *P* (**L**), where *P* (**L**) contains the loading penalties described below. This is equivalent to maximum a posteriori estimation of the latent factor activities under their Brownian priors. Brownian densities and gradients were evaluated by Gaussian pruning along the tree, avoiding repeated construction or inversion of dense covariance matrices.

Loading entries were initialized from a standard normal distribution and normalized by row. A provisional **F** = **XL**^T^ was used only to initialize the Brownian variance parameters. For each provisional factor, its empirical variance across cells was used to choose an initial 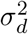 that produced the same expected variance among the tree tips. Before each objective evaluation, including the first, **F** was set to its exact conditional optimum given the current loadings and variance parameters. This update diagonalized **Σ** once and solved a *k* × *k* linear system for each tree eigenmode. Each entry of the loading matrix **L** and the logarithms of the variance parameters were optimized as free parameters using Adam [79] with a learning rate of 0.01 and adaptive gradient-norm clipping. Brownian variances were bounded below by 10^−6^, and additional loading constraints were enforced as described below.

Optimization proceeded for at least 500 iterations. Convergence was assessed from the mean objective over successive 50-iteration windows and declared when the relative improvement was no greater than 10^−6^. The parameter state with the lowest observed objective was retained. The factor dimension *k* must be specified and is then fixed within each fit.

### Lineage-based gene expression simulation

We simulated ultrametric cell-lineage trees using the Cassiopeia (v2.1.0) [80] birth–death fitness simulator, with birth and death rates of 0.075 and 0.005, respectively, and rescaled each tree to unit height. Gene expression and latent factors were then generated forward under the same Brownian model used for inference. For a tree with covariance matrix **Σ**, each latent factor was drawn as

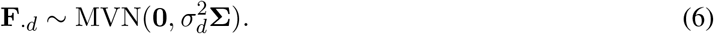

To evaluate phylogenetic gene filtering, we simulated 10 trees at each of 25, 50, 100, 500, 1,000, 5,000, and 10,000 cells. For every tree, we generated 1,000 heritable genes under Brownian motion and 1,000 non-heritable genes using identity covariance, with both sets having variance 0.25 at the tips. These known classes were used to calculate the precision and recall of each filtering method.

To evaluate factor-model recovery, we simulated 10 datasets for each combination of 500, 1,000, or 5,000 cells and 500, 1,000, or 5,000 genes. Each dataset contained five factors with Brownian variances (20, 10, 5, 2, 1). Loading entries were drawn independently from a standard normal distribution and each loading row was rescaled to unit Euclidean norm. We generated the observed expression matrix as

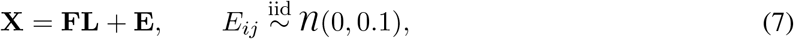

and fit the model directly to these simulated matrices without phylogenetic gene filtering, allowing the inferred and generating factors, loadings, covariance structures, and objective components to be compared directly.

### Statistical tests for phylogenetic gene filtering

For the provided tree, we calculated the Brownian covariance matrix **Σ**, whose entry Σ_*ij*_ is the shared root-to-ancestor time for cells *i* and *j*. Let **y**_*g*_ denote the centered expression of gene *g* across *n* cells.

For the autocorrelation test, we followed the approach introduced in PATH (v1.0) [61] and confirmed that our implementation was numerically identical to the original. Pairwise patristic distances were calculated as *d*_*ij*_ = Σ_*ii*_ + Σ_*jj*_ − 2Σ_*ij*_ and converted to weights *w*_*ij*_ = 1*/d*_*ij*_ for *i* ≠ *j*, with *w*_*ii*_ = 0. The full weight matrix was then normalized so that Σ_*ij*_ *w*_*ij*_ = 1. Phylogenetic autocorrelation was measured using Moran’s statistic,

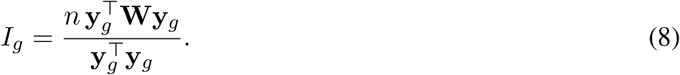

We compared the observed *I*_*g*_ with its expected value under random assignment of expression values across cells, *E*[*I*_*g*_] = −1*/*(*n* − 1). We divided this difference by its analytic standard deviation, as defined in PATH, to obtain a *z* score and calculated a two-sided normal *P* value.

The full Brownian likelihood-ratio test compared an independent Gaussian null model, 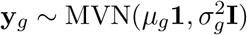, with the Brownian alternative 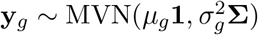. Under each model, the mean was estimated by generalized least squares and the variance by its closed-form maximum-likelihood estimate, thereby profiling both parameters without numerical optimization. The test statistic was

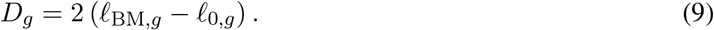

Because these covariance models are not nested through a single fitted parameter, we generated 1,000 independent Gaussian datasets under the gene-specific null model and calculated the *P* value as the fraction producing a statistic at least as large as *D*_*g*_.

The Pagel’s *λ* likelihood-ratio test provided a nested version of this comparison, following its original phylogenetic formulation [75, 76] and our previous application to lineage-traced expression data [40]. For each gene, we formed **Σ**(*λ*) by retaining the diagonal of **Σ** and multiplying its off-diagonal entries by *λ*, then maximized the profiled Gaussian likelihood over 0 ≤ *λ* ≤ 1 using bounded Brent optimization, with both boundary values also evaluated explicitly. Thus, *λ* = 0 represents phylogenetically independent expression and increasing *λ* restores the covariance expected from the lineage tree. We calculated 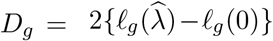 and obtained its *P* value from the 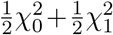 boundary-null distribution. For all three tests, *P* values were adjusted across genes using the Benjamini–Hochberg procedure, and genes with *q* ≤ 0.05 were retained. The likelihood-ratio tests additionally required *D*_*g*_ *>* 0.

### Factor-model identifiability constraints

The factorization **X ≈ FL** is unchanged if a column **F**_·*d*_ is multiplied by a constant and the corresponding loading row **L**_*d*·_ is divided by the same constant. Paired sign changes and factor permutations are likewise equivalent. We fixed the scale of each factor by constraining every row of the loading matrix to have unit Euclidean norm,

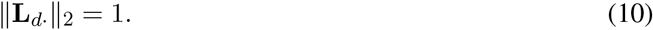

During optimization, loading gradients were projected onto the tangent space of this constraint and loading rows were renormalized after each update. The corresponding column of **F** therefore contained the factor’s fitted scale and variation across cells.

After fitting, factors were ordered by decreasing Brownian variance 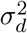. Because simultaneously multiplying **F**_·*d*_ and **L**_*d*·_ by −1 leaves their product unchanged, we also oriented each factor so that Σ_*j*_ *L*_*dj*_ ≥ 0, reversing both its scores and loadings when necessary. These conventions do not change the fitted expression matrix, but provide consistent factor scales, labels, and directions for reporting and comparison.

### Factor-loading regularization

We regularized the factor-loading matrix **L** by adding *ℓ*_1_ and loading-overlap penalties to the model objective. For *n* cells, *p* genes, and *k* factors, the combined penalty was

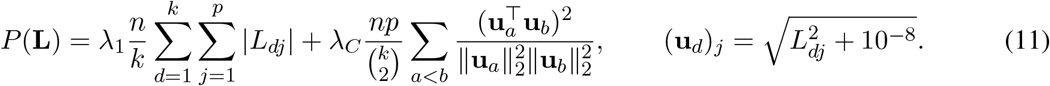

The *ℓ*_1_ term favors factors defined by a sparse subset of genes. The second term is the squared cosine similarity between the absolute loading magnitudes of each factor pair and therefore discourages different factors from repeatedly using the same genes, regardless of loading sign. The scaling factors account for the size of the expression matrix and the number of factors or factor pairs in the objective.

### Data acquisition, preprocessing, and VINE lineage reconstruction

We obtained paired single-cell RNA sequencing and CRISPR lineage-recording data for the TLS1 and TLS2 mouse trunk-like structures from GSE220949 [62], and for the CP1–CP4 lineage groups from the mouse 5k lung-cancer xenograft experiment in GSE161363 [10]. For the developmental data, allele tables were converted to integer character matrices using Cassiopeia, after which characters and cells containing only unedited or missing states were removed. For the cancer data, we used the deposited allele-threshold character matrices and encoded missing states as −1. In both datasets, the 10x count matrix was restricted and reordered to the cells represented in the corresponding character matrix, and genes with no counts among the retained cells per clone were removed.

Lineage trees were inferred separately for each sample or clone using VINE (v0.3.5) [81] with its CRISPR model. We required at least 400 variational iterations and assessed convergence over a window of 100 iterations. Each run produced 100 approximate posterior tree samples. The first sampled tree from each run was used in the present analyses.

### Gene set enrichment and Gene Ontology analysis procedures

Gene Ontology (GO) and gene set enrichment analyses were performed with clusterProfiler (v4.10.0) [82] in R (v4.3.3). Gene symbols were mapped to Entrez identifiers with `bitr` using org.Mm.eg.db or org.Hs.eg.db (v3.18.0) for the mouse and human datasets, respectively. To characterize the binary gene-filtering results, genes passing and not passing the phylogenetic filter were tested separately for over-representation of GO Biological Process terms using `enrichGO`, with all mapped genes tested in that sample used as the background. To characterize the continuous factor loadings, mapped genes were ranked from the most positive to the most negative loading for each factor and tested for GO Biological Process enrichment using `gseGO`. Zero and nonfinite loadings were excluded, duplicate Entrez identifiers were represented by the loading with the largest absolute value, and the sign of the normalized enrichment score identified enrichment toward the positive or negative loading direction.

### Latent-factor ancestral-state reconstruction

For each fitted factor *d*, we reconstructed its value at every internal node conditional on the fitted tip scores **F**_·*d*_. Under the Brownian model, information about a node can be represented by a Gaussian summary (*µ, v*), where *µ* is the estimated state and *v* is its uncertainty. Transporting this summary across a branch of length *t* leaves its mean unchanged and increases its variance from *v* to 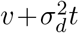. Two independent summaries (*µ*_1_, *v*_1_) and (*µ*_2_, *v*_2_) for the same node were combined by precision weighting,

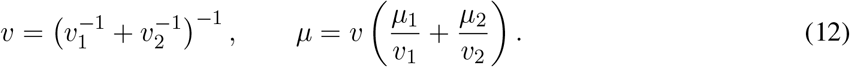

Zero variance denotes a fixed state and was handled as the limiting case of this calculation.

We applied this calculation in two tree traversals. First, a postorder traversal summarized the information below each node. Each tip began with the fixed state (*F*_*id*_, 0). The summaries from its two children were transported across their respective branches and combined at the parent. Second, a preorder traversal summarized the complementary information arriving from above each node. This traversal began at the root with the Brownian prior 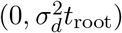. The summary passed to a child was obtained by combining the summary from above its parent with the transported summary from its sibling, and then transporting the result across the branch to the child. Finally, the descendant and complementary summaries were combined at each internal node. The resulting mean *F*_*id*_ is therefore conditional on all fitted tip states, while tip values remain equal to their fitted factor scores.

### Hotspot gene-module inference

We identified lineage-associated gene modules with Hotspot (v1.1.3) [51], following the parameter settings in its lineage-data tutorial (https://hotspot.readthedocs.io/en/latest/Lineage_Tutorial.html). For each clone, we retained cells present in both the count matrix and lineage tree, pruned the tree to those cells while preserving branch lengths, and retained genes detected in at least 10 cells. Because raw counts were available, we used Hotspot’s depth-adjusted negative-binomial model (`danb`) with each cell’s total UMI count as its scaling factor. We constructed an unweighted lineage-neighborhood graph with 30 neighbors. We first calculated gene-level autocorrelation and retained genes with FDR *<* 0.05. Pairwise local correlations were then calculated among all retained genes. Modules were obtained by agglomerative clustering with a minimum module size of 50 genes, an FDR threshold of 0.05 for merging branches, and `core_only=True` so that genes with ambiguous module membership were left unassigned. Per-cell module scores were calculated using `hs.calculate_module_scores()`. For visualization along the lineage tree, the score at each internal node was defined as the mean score among its descendant cells.

### Pearson correlation analysis of factor loadings and Hotspot modules

We compared fitted factors across clones using their complete gene-loading profiles rather than only their top genes. Factors with Brownian variance 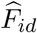 were retained as active, and the two loading matrices were restricted to the intersection of genes present in both fits. For factors *a* and *b*, with loadings *L*_*ag*_ and *L*_*bg*_ over the shared gene set *G*, we calculated

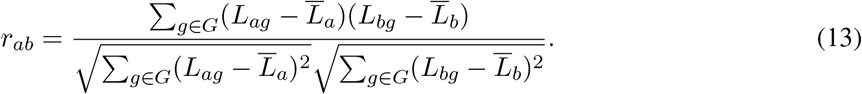

No additional scaling or imputation was applied before calculating these correlations.

To compare factors with Hotspot, we used the intersection of genes in the factor-loading matrix and the full set of genes tested for Hotspot autocorrelation. For each Hotspot module *m*, we defined a binary vector *H*_*mg*_ equal to one when gene *g* was assigned to *m* and zero when it was assigned to another module or remained unassigned. Pearson correlation between *H*_*m*·_ and each active factor-loading vector *L*_*a*·_ then measured whether genes assigned to a module tended to have consistently positive or negative loadings on that factor. This is equivalent to a point-biserial correlation. For visualization only, rows and columns of each correlation matrix were ordered using a maximum-weight one-to-one assignment based on absolute correlation. This ordering did not alter the reported correlation values.

### Cross-context comparison of factor or module biological functions

To compare factors between the developmental and lung cancer datasets, we collected GO biological process terms with GSEA-adjusted *P* ≤ 0.05 across both loading directions. For a developmental factor *a* and cancer factor *b*, with significant term sets *S*_*a*_ and *S*_*b*_, we calculated the unsigned Jaccard similarity

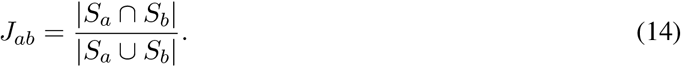

Pairs with no terms in their union were assigned a similarity of zero. For visualization only, rows and columns were ordered using a maximum-weight one-to-one assignment based on Jaccard similarity, which did not alter the similarity values.

We applied the same procedure to the existing GO over-representation results for Hotspot modules. Modules with at least one GO biological process term having adjusted *P* ≤ 0.05 were retained, and Jaccard similarities were calculated between the significant term sets of each developmental and cancer module.

### Software implementation

The phylogenetic signal gene filtering models, phylogenetic factor analysis model, and simulation tool were all implemented in C (C99). Source code is available at https://github.com/StephenStaklinski/scPFA under the BSD 3-Clause License. Any additional pipelines and scripts needed to reproduce the analyses presented here are available at https://github.com/StephenStaklinski/scPFA_analyses.

## Acknowledgments

We thank other members of the community at Cold Spring Harbor Laboratory for helpful feedback, particularly Bruce Stillman, David M. McCandlish, and Hannah V. Meyer.

