## Supplementary Material for "Lineage-informed factor analysis reveals heritable programs of single-cell gene expression"

### Supplementary Information for:
### Lineage-informed factor analysis reveals heritable programs of single-cell gene expression

Stephen J. Staklinski[1] and Adam Siepel[1,*]

[1]Simons Center for Quantitative Biology, Cold Spring Harbor Laboratory, Cold Spring Harbor, NY

#### Supplementary figures

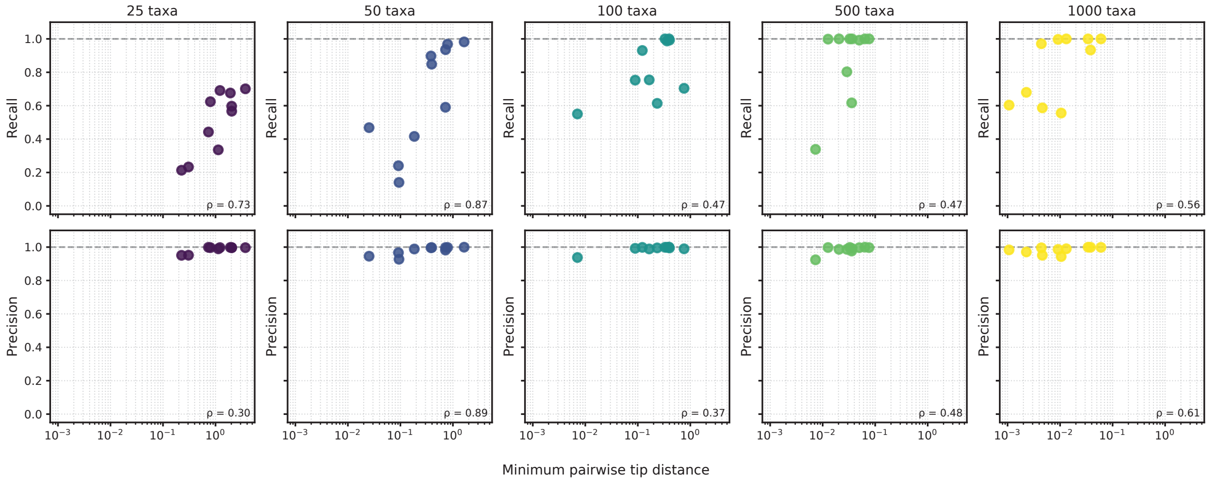


**Supplementary Figure S1: Minimum pairwise tip distance is associated with the performance of the PATH phylogenetic gene filter.** Precision (bottom row) and recall (top row) of the PATH phylogenetic gene filter are plotted against the minimum pairwise tip distance of the simulated lineage tree for datasets of varying size (columns). Each point represents a simulation replicate, and $\rho$ denotes the Pearson correlation coefficient.

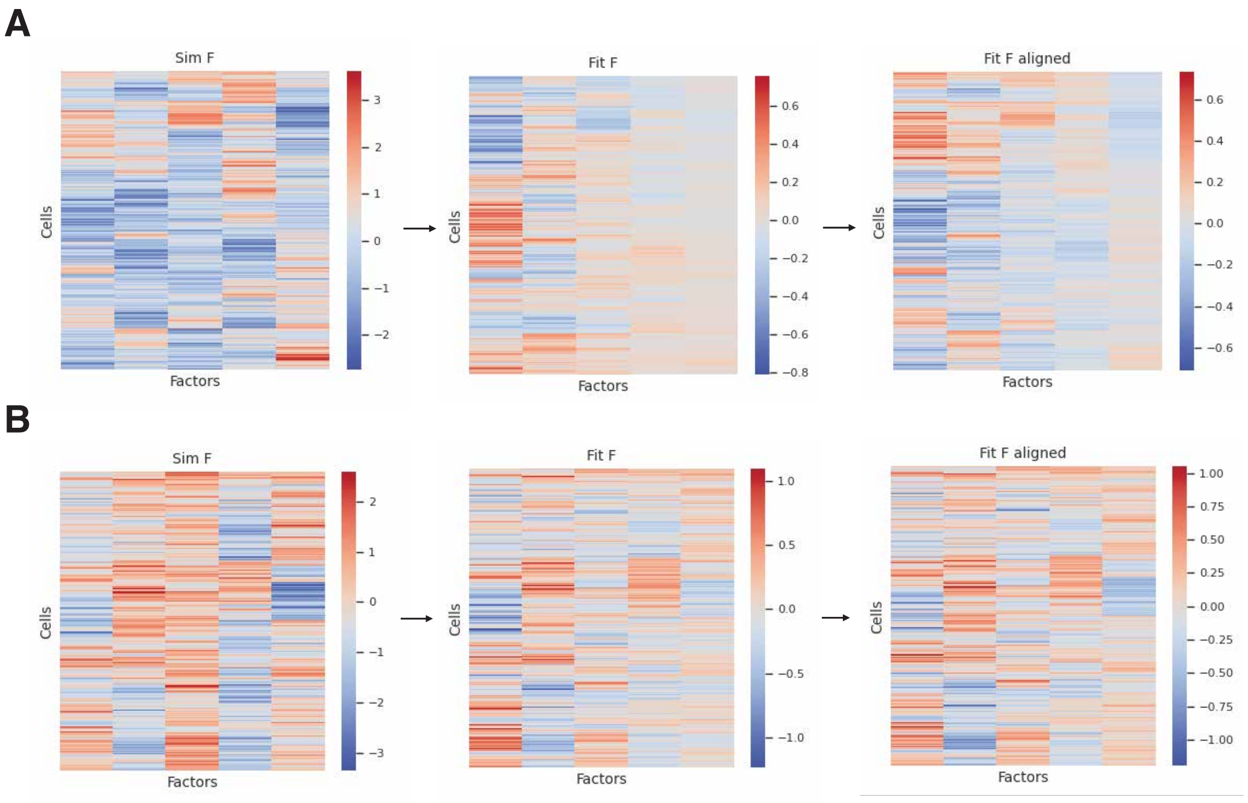


**Supplementary Figure S2: Example recovery of latent factors before and after identifiability constraints.** **(A)** Simulated ground-truth cell-factor matrix (**F**) (left), the corresponding unconstrained model fit (middle), and the fit after Procrustes alignment (right). **(B)** Same as (A), but with scale, permutation, and sign constraints applied during model fitting.

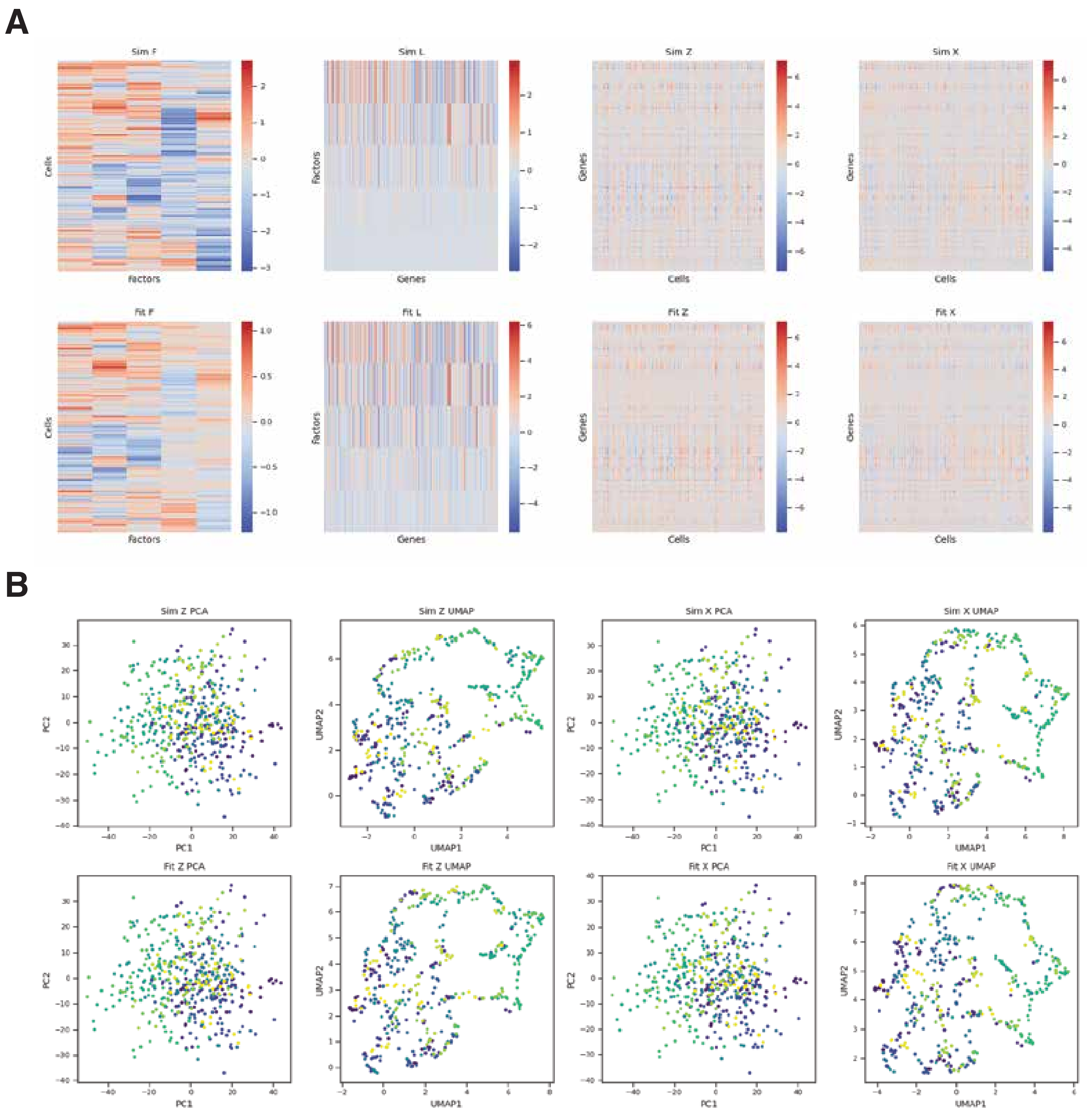


**Supplementary Figure S3: Representative simulation demonstrating recovery of latent factors and reconstructed gene expression.** **(A)** Example simulation with 500 cells and 500 genes. Columns show the cell-factor matrix ($\mathbf{F}$), factor-gene loading matrix ($\mathbf{L}$), reconstructed latent gene expression matrix ($\mathbf{Z} = \mathbf{FL}$), and observed gene expression matrix ($\mathbf{X}$). The top row shows the simulated ground truth and the bottom row shows the corresponding model fit. **(B)** Low-dimensional representations of simulated (top) and fitted (bottom) gene expression matrices. The first two columns show PCA and UMAP projections of the latent expression matrix ($\mathbf{Z}$), while the final two columns show PCA and UMAP projections of the observed expression matrix ($\mathbf{X}$). Each point corresponds to a cell and is colored arbitrarily in the matched order of cells in the matrix to enable relative comparisons.

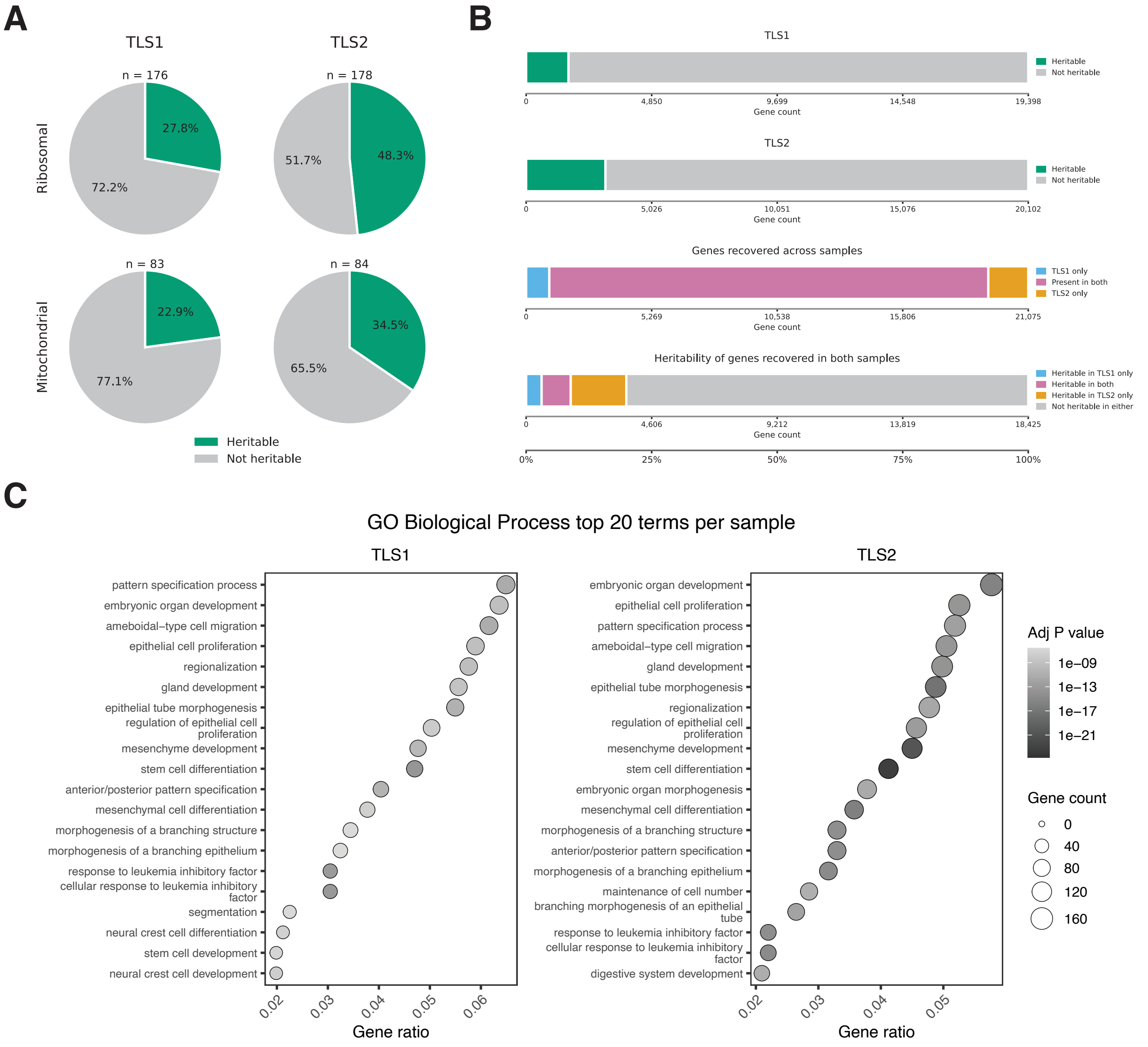


**Supplementary Figure S4: Genes with phylogenetic signal capture shared developmental processes.** **(A)** Fractions of ribosomal and mitochondrial genes with phylogenetic signal in TLS1 and TLS2. **(B)** Fractions of all genes with phylogenetic signal in each clone (top), overlap in genes recovered across clones (middle), and clone specificity of phylogenetic signal among shared genes (bottom). **(C)** Top 20 enriched Gene Ontology (GO) biological processes among genes with phylogenetic signal in TLS1 and TLS2. Point size indicates gene count, color indicates adjusted $P$ value, and the horizontal axis shows the fraction of genes associated with each term.

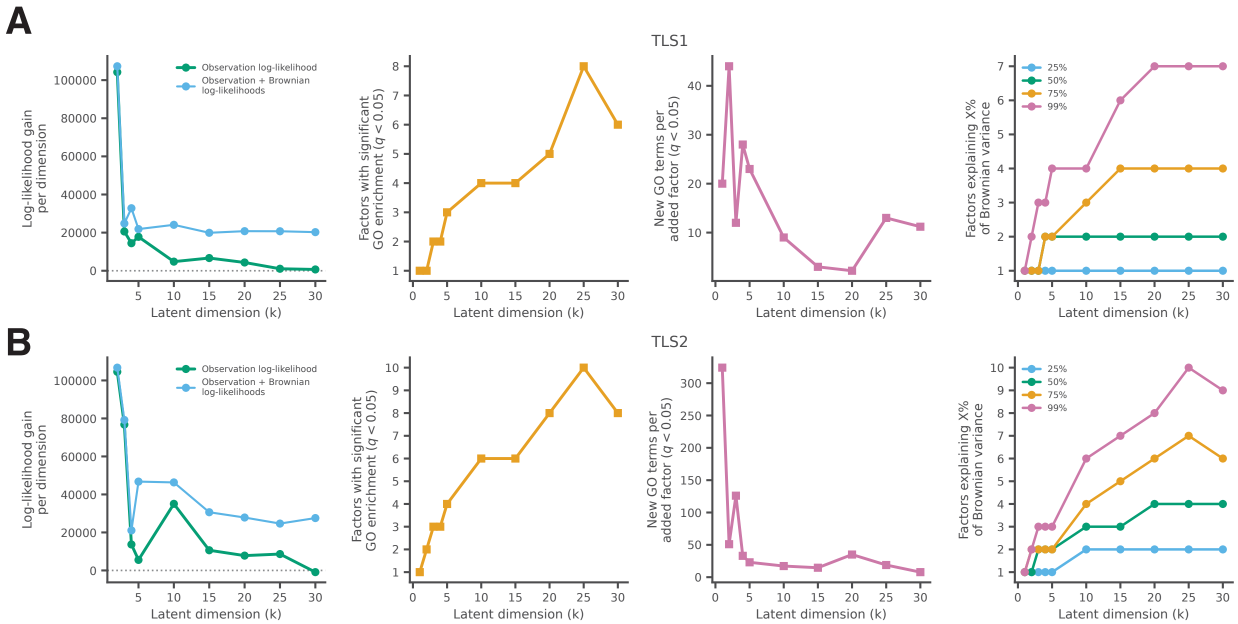


**Supplementary Figure S5: Dimensionality selection for developmental lineage-tracing datasets. (A–B)** Model diagnostics across latent dimensions $k$ for TLS1 (A) and TLS2 (B). From left to right: gain in observation and combined observation–Brownian log-likelihood per added dimension; number of factors with significant GO enrichment ($q < 0.05$); number of new significant GO terms per added factor; and number of factors required to explain 25%, 50%, 75%, and 99% of the total Brownian variance.

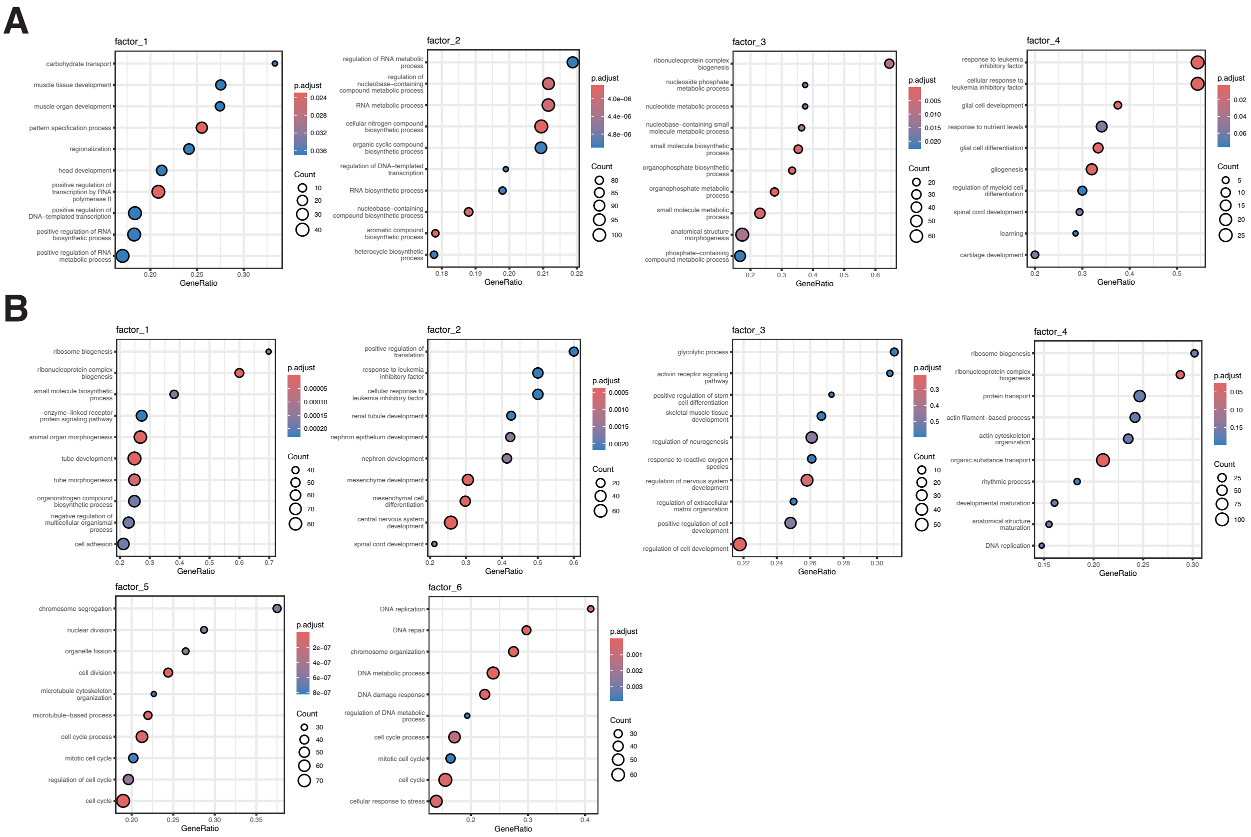


**Supplementary Figure S6: Developmental factors are enriched for distinct biological processes. (A–B)** Gene Ontology biological process enrichment for active factors in TLS1 (A) and TLS2 (B). Point size indicates gene count, color indicates adjusted $P$ value, and the horizontal axis shows the fraction of genes associated with each term.

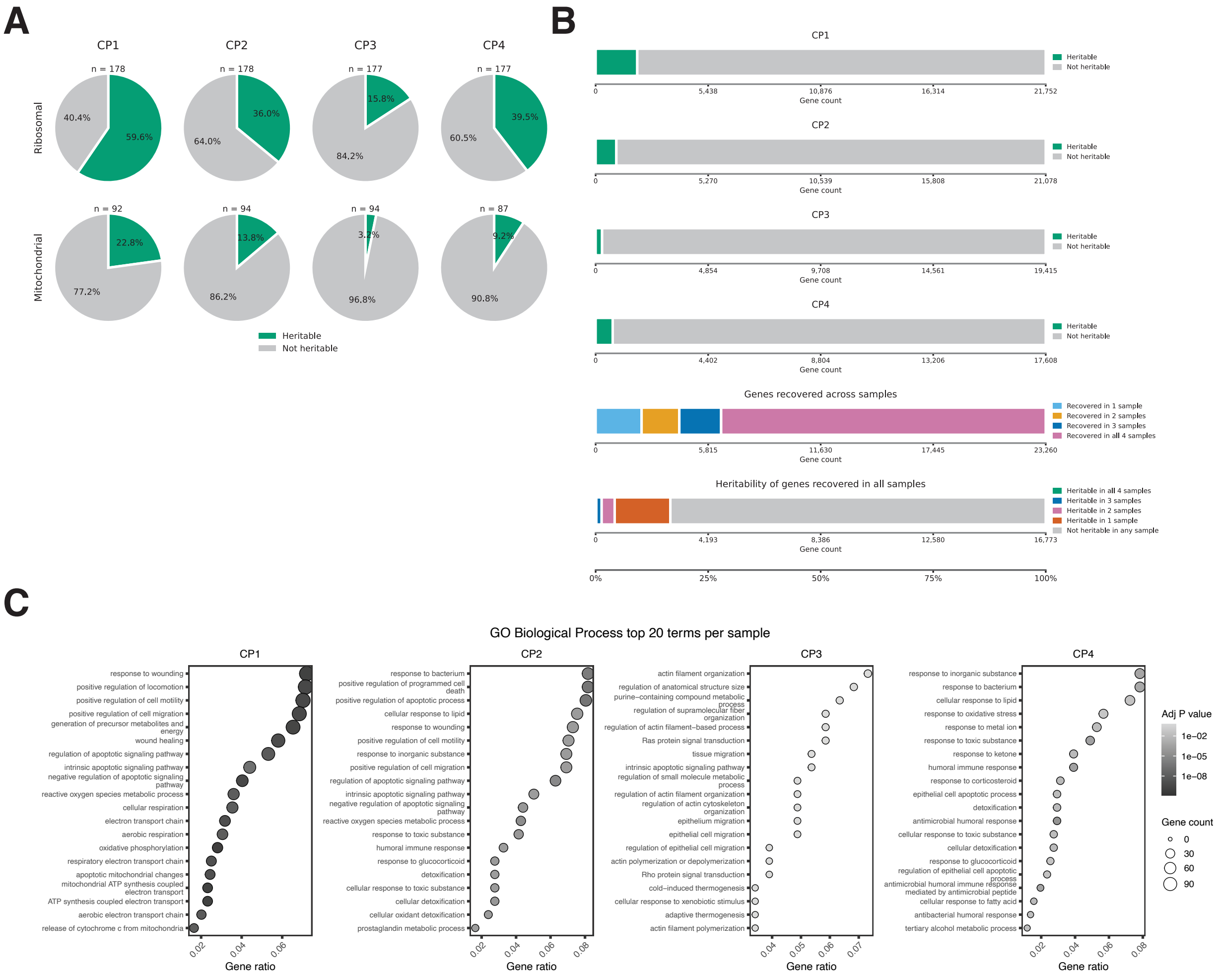


**Supplementary Figure S7: Genes with phylogenetic signal capture biological processes across cancer clones.** **(A)** Fractions of ribosomal and mitochondrial genes with phylogenetic signal in clones CP1–CP4. **(B)** Fractions of all genes with phylogenetic signal in each clone (top), number of clones in which each gene was recovered (middle), and number of clones in which shared genes showed phylogenetic signal (bottom). **(C)** Top 20 enriched Gene Ontology (GO) biological processes among genes with phylogenetic signal in each clone. Point size indicates gene count, color indicates adjusted $P$ value, and the horizontal axis shows the fraction of genes associated with each term.

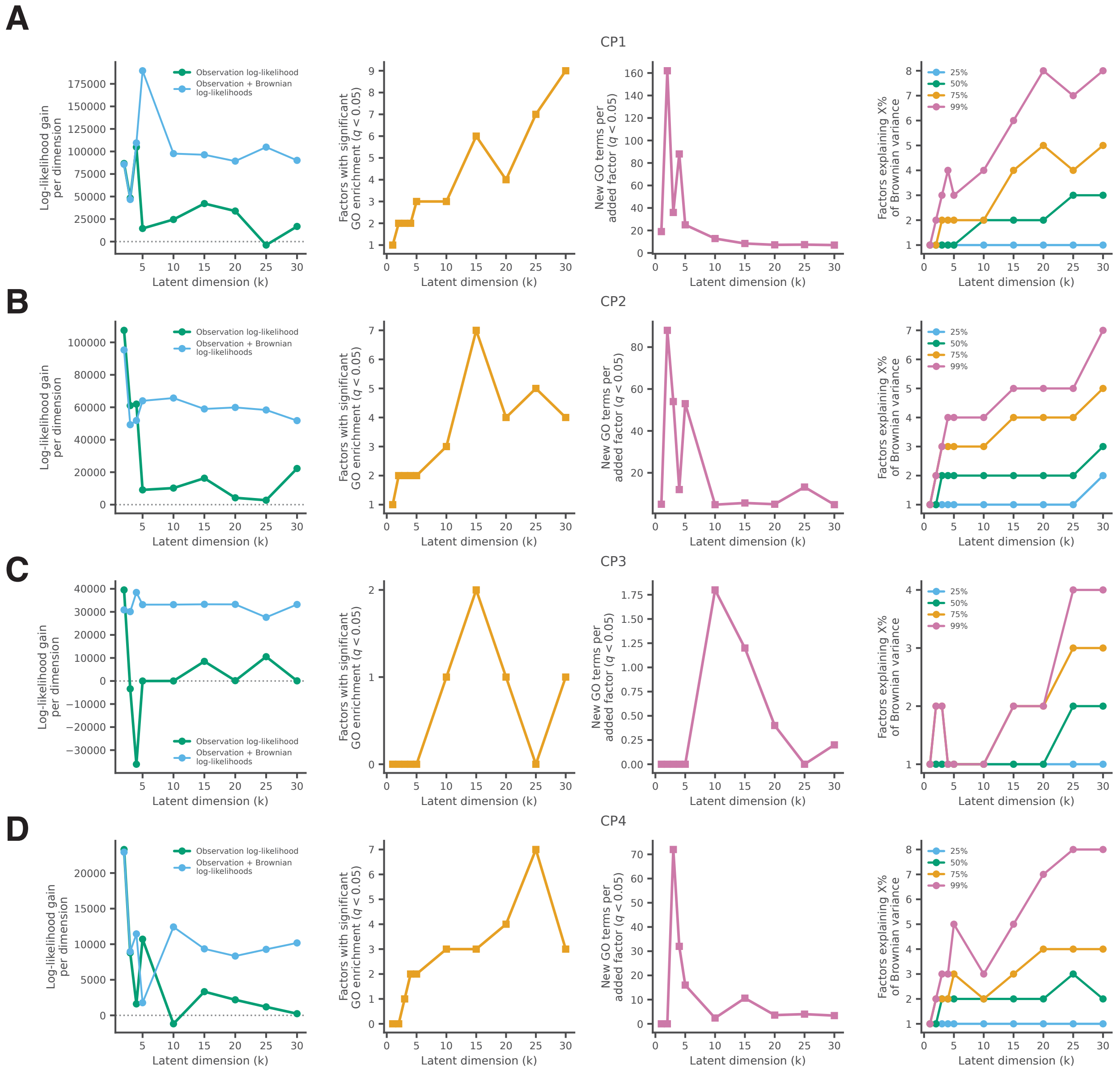


**Supplementary Figure S8: Dimensionality selection for lung cancer lineage-tracing datasets. (A–D)** Model diagnostics across latent dimensions $k$ for CP1–CP4, respectively. From left to right: gain in observation and combined observation–Brownian log-likelihood per added dimension; number of factors with significant GO enrichment ($q < 0.05$); number of new significant GO terms per added factor; and number of factors required to explain 25%, 50%, 75%, and 99% of the total Brownian variance.

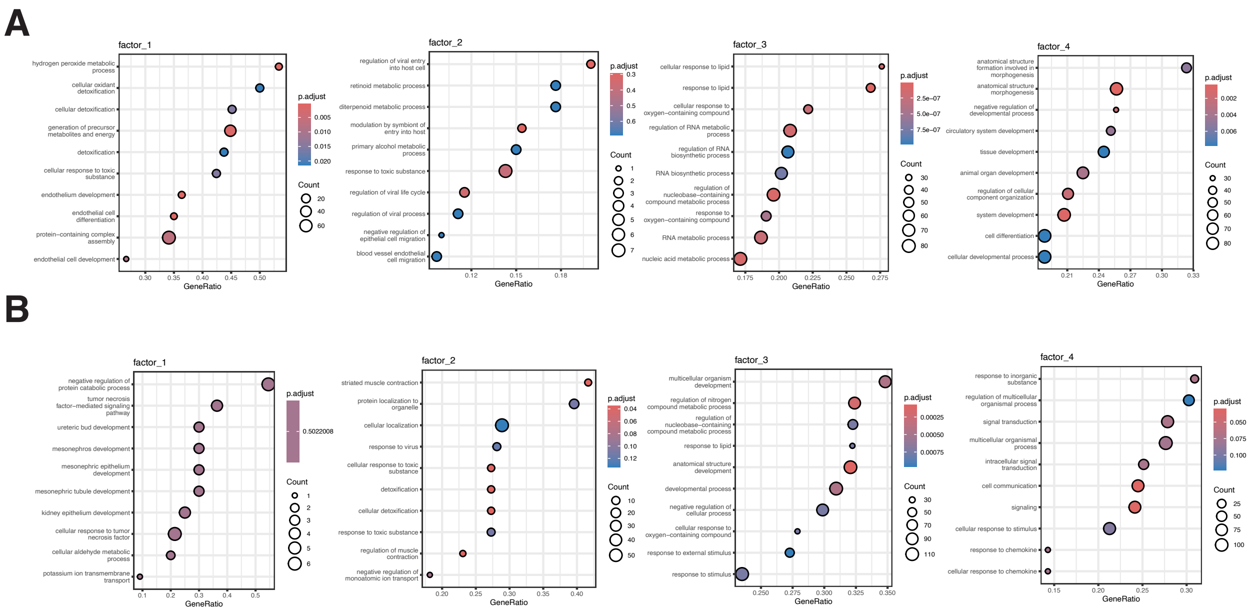


**Supplementary Figure S9: Lung cancer factors are enriched for distinct biological processes. (A–B)** Gene Ontology biological process enrichment for active factors in CP1 (A) and CP2 (B). Point size indicates gene count, color indicates adjusted $P$ value, and the horizontal axis shows the fraction of genes associated with each term.

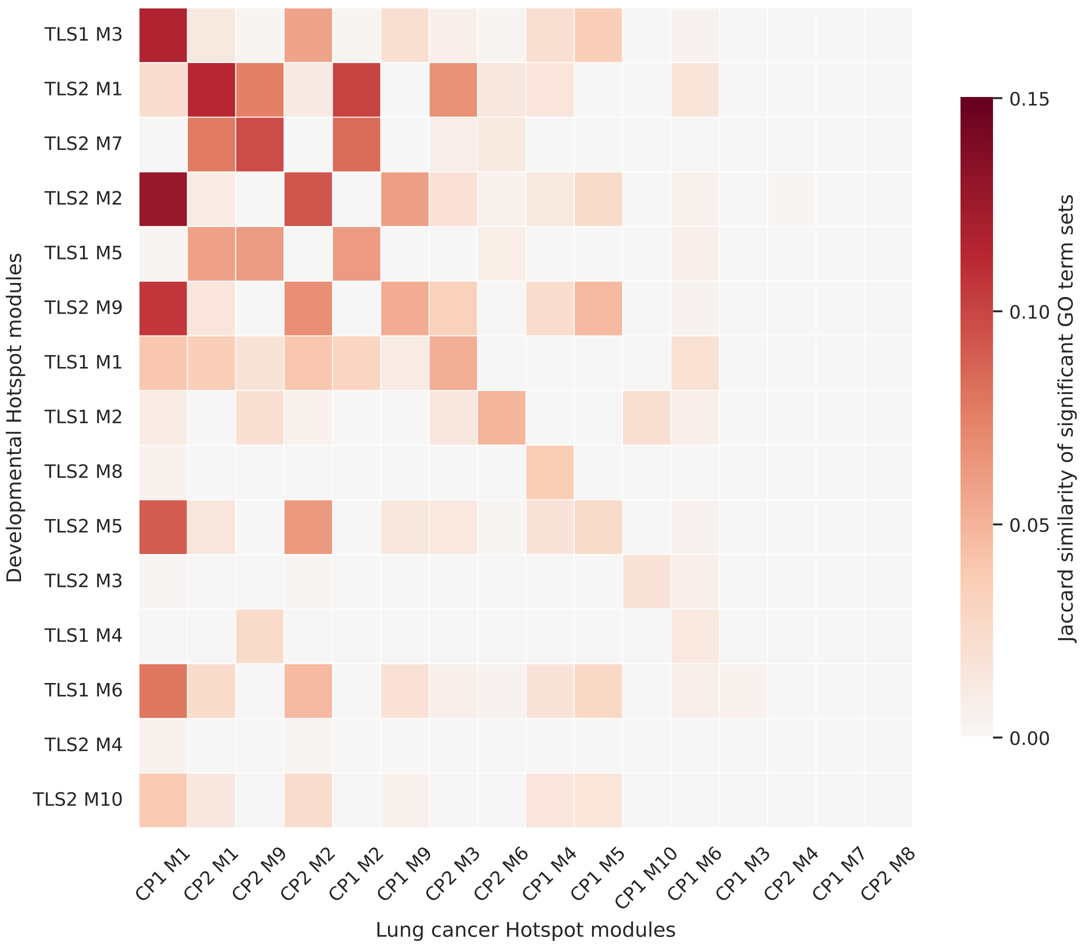


**Supplementary Figure S10: Hotspot modules are functionally distinct across developmental and lung cancer systems.** Jaccard similarity between sets of significantly enriched Gene Ontology biological process terms for developmental Hotspot modules (rows) and lung cancer Hotspot modules (columns). Only modules with at least one significant term (adjusted $P \leq 0.05$) are shown. Rows and columns are ordered around the strongest one-to-one matches.
